# Structural basis of *γ*-chain family receptor sharing at the membrane level

**DOI:** 10.1101/2023.05.05.539662

**Authors:** Tiantian Cai, Rachel Lenoir Capello, Xiong Pi, Hao Wu, James J. Chou

## Abstract

The common γ-chain (γc) family of cytokine receptors, including interleukin (IL)-2, IL-4, IL-7, IL-9, IL-15, and IL-21 receptors, are activated upon engagement with the common γc receptor in ligand dependent manner. Sharing of γc by the IL receptors (ILRs) is thought to be achieved by concomitant binding of γc and ILR ectodomains to a cytokine. Here, we found that direct interactions between the transmembrane domain (TMD) of γc and those of the ILRs are also required for receptor activation, and remarkably, the same γc TMD can specifically recognize multiple ILR TMDs of diverse sequences. Heterodimer structures of γc TMD bound to the TMDs of IL-7R and IL-9R, determined in near lipid bilayer environment, reveal a conserved knob-into-hole mechanism of recognition that mediates receptor sharing within the membrane. Functional mutagenesis data indicate the requirement of the heterotypic interactions of TMDs in signaling, which could explain disease mutations within the receptor TMDs.

**One-Sentence Summary:** The transmembrane anchors of interleukin receptors of the gamma-chain family are critical for receptor sharing and activation.

---

Members of the common γ-chain family of cytokine receptors, as the name indicates, have the feature that upon binding to their matching cytokines, they pair specifically with a common, shared γ-chain (γc) to allow signal transduction (*1, 2*). To date, the family consists of six members, including the interleukin-2 receptor (IL-2R) from which the γc was first discovered (*3*) and other interleukin receptors (ILRs) with specificity for IL-4, IL-7, IL-9, IL-15, and IL-21, respectively. These receptors play crucial roles in the development and proliferation of multiple lymphocyte lineages of both innate and adaptive immune systems (*1, 4*), and are thus clinically important targets for immunotherapy (*2, 5, 6*). ILRs and γc are typical Type-I transmembrane proteins comprising a ligand-binding ectodomain, a transmembrane domain (TMD), and an intracellular domain (ICD) that contains the Janus kinase (JAK) binding sites. In addition to the γc receptor family, there are two other major families of class I cytokine receptors that share common receptors: one uses the common gp130 receptor (*7*) and the other shares the β-chain (βc) receptor (*8*).

The mechanism by which cytokines or interleukins mediate specific pairing of the γc with ILRs has been unraveled by crystallographic studies of the IL-2-bound and IL-4-bound receptor complexes (*9-11*). In both cases, the cytokine binding to an ILR generates a composite surface for subsequent interaction with the γc. Although the interleukins recognized by the γc have ∼15% sequence identity among them, the γc mostly interacts with the last helix (Helix D) of the 4-helix bundle interleukin structures and Helix D is much more conserved (∼35% identity). Notably, this interaction interface is small (∼1000Å^2^) and is characterized by relatively flat surface complementarity (*10*). These structural features constitute the structural basis for the degenerate cytokine recognition by the γc.

An aspect of ILRs not well understood is the TMD and its possible role in receptor signaling, and this question was inspired by previous studies on other receptors such as the epidermal growth factor receptor (EGFR) and receptors in the tumor necrosis factor receptor superfamily (TNFRSF) that revealed much more active roles of TMDs in mediating receptor oligomerization and activation than conventionally appreciated (*12-19*). An early cryo-EM study of a full-length, transmembrane form of a quaternary cytokine-receptor complex consisting of gp130, LIF-R, the cytokine Ciliary Neurotrophic Factor (CNTF), and its alpha receptor (CNTF-Rα) suggests that the TMDs of these receptor chains are associated (*20*). More importantly, for the γc family receptors, both gain- and loss-of-function disease mutations within the TMD have been reported, e.g., mutations or frameshifts within the core of the γc TMD have been linked to X-linked severe combined immunodeficiency diseases (X-SCID) (*21-23*), but the structural basis of these disease mutations is unknown. Thus far, no structural information is available for the transmembrane region of any of the γc family receptors.

In this study, we found that heterotypic interactions between the TMDs of the γc and its family members are an essential component of receptor signaling, raising the question of how a single transmembrane helix (TMH) can have the structural sophistication to specifically recognize a variety of TMHs with highly divergent sequences. We then determined high resolution structures of the γc TMD in complex with IL-7R and IL-9R TMDs, respectively, in bicelles that mimic a lipid bilayer. The two heterodimer structures and extensive functional mutagenesis reveal a common knob-into-hole mechanism that underlies degenerate ILR pairing with the γc within the membrane.

## RESULTS

### Removal of the ectodomains of IL-7R and γc activates IL-7R signaling

A large body of structural and functional studies on members of the γc family receptors have painted an overall picture of receptor activation (*5, 9-11, 24*). Briefly, interleukins bind to the ectodomains of their α or β chain receptors to form a complex that subsequently recruits the γc, thus positioning the ICDs in the right arrangement to allow reciprocal phosphorylation of JAK1 and JAK3 and activate downstream JAK/STAT signaling (Fig. 1A). The ectodomain and ICD are, however, separated by the TMD, which could also play a role in driving the JAKs into signaling-compatible configuration as they contain loss-of-function disease mutations (*21-23*). In this case, the receptor ectodomains before ligand engagement should physically prevent ligand-independent, TMD-driven JAK1/JAK3 phosphorylation. To test whether removal of ectodomains can free the TMDs to mediate heterotypic association between ILR and γc, we inserted a TEV cleavage site (ENLYFQGGGGGS) between the ectodomain and TMD for both human IL-7R (TEV-hIL-7R) and γc (TEV-hγc) as illustrated in Fig. 1B. BaF3 cells with co-expressed wild-type (WT) or TEV-inserted receptors were incubated with TEV protease to shed the ectodomains, and JAK/STAT activation was detected by immunoblot analysis of STAT5 phosphorylation.

**Fig. 1.**
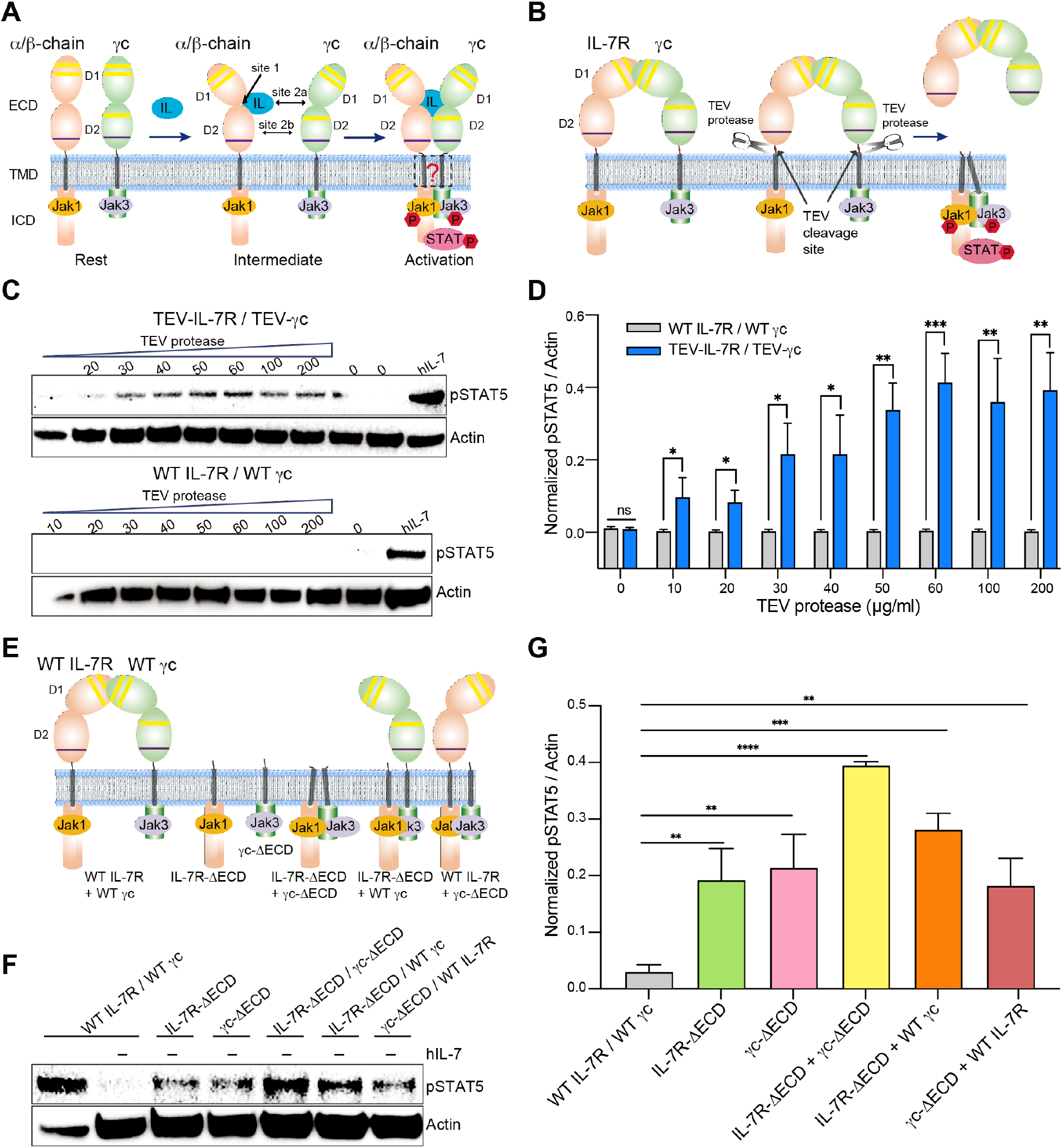
Removal of the ectodomains of IL-7R and γc activates IL-7R signaling. (**A**) The prevailing model of cytokine mediated receptor sharing for receptors in the common γc receptor family. Receptors in this family are Type I membrane proteins comprising a ligand binding ectodomain (ECD), a transmembrane domain (TMD), and an intracellular domain (ICD) with docking sites for the JAK/STAT signaling molecules. (**B**) The autoinhibition hypothesis postulating that preligand association of ECDs hinders clustering of TMD-ICD for activation and that proteolytic removal of the ECDs should allow TMD-driven receptor pairing and cytokine-independent signaling. (**C**) IL-7R pathway activation detected by immunoblotting of STAT5 phosphorylation (pSTAT5) in BaF3 cells co-expressing cleavable receptors (TEV-IL-7R / TEV-γc, upper panel) or WT receptors (WT IL-7R / WT γc, lower panel), after treatment with 0-200 μg/ml TEV protease or 50 ng/ml hIL-7. (**D**) Quantification of protease-induced pSTAT5 signals in (C) by ImageJ as pSTAT5/Actin intensity ratios, which are further normalized relative to that induced by IL-7. Results are from 3 independent experiments (n = 3) and expressed as mean ± SEM. Statistical significance: t-test, ns, not significant; *P≤0.05; **P≤0.01; ***P≤0.001. (**E**) Schematic illustration of all possible combinations of full-length receptors (IL-7R, γc) and ECD-deleted receptors (IL-7R-ΔECD, γc-ΔECD) expressed in BaF3 cells for testing ligand independent signaling. (**F**) IL-7R pathway activation reported by STAT5 phosphorylation in BaF3 cells expressing full-length and ECD-deleted receptor combinations illustrated in (E). (**G**) Quantification of pSTAT5 signals in (F) by ImageJ as pSTAT5/Actin intensity ratios, which are further normalized relative to that induced by IL-7. Results are from 3 independent experiments (n = 3) and expressed as mean ± SEM. Statistical significance: t-test; ns, not significant; *P≤0.05; **P≤0.01; ***P≤0.001; ****P < 0.0001.

Both WT receptors (hIL-7R, hγc) and cleavable receptors (TEV-hIL-7R, TEV-hγc) expressed well on the cell membrane with EGFP and mCherry fused to the C-termini of hIL-7R and hγc, respectively (fig. S1A). Co-localization of hIL-7R and hγc, as well as sparse bright puncta, were observed in the absence of hIL-7, suggesting that these receptors may pre-associate even in the absence of ligand. More puncta appeared on the cell surface after ligand incubation due to greater receptor clustering induced by ligand binding, and expectedly, this was accompanied by receptor activation as indicated by STAT5 phosphorylation (fig. S1B). The TEV-cleavable hIL-7R and hγc exhibited very similar ligand response as the WT receptors (fig. S1B) but, remarkably, could be activated by incubating BaF3 cells with TEV protease in the absence of ligand (Fig. 1C). TEV protease dose-response profile indicates that ∼40% maximal activation can be reached compared to IL-7 induced activation (Fig. 1D). Accordingly, more puncta on the cell surface were observed after treatment with 50μg/ml TEV protease, suggesting the ability of TMH-ICD to cluster though significantly weaker than in the context of ectodomain-ligand engagement (fig. S1C). In contrast, no activation was detected after adding TEV protease to cells expressing the WT receptors (Fig. 1, C to D and fig. S1C).

To independently examine the functional consequence of ectodomain removal, we designed two constructs with the ectodomain deleted (hIL-7R-ΔECD and hγc-ΔECD) and expressed each of them alone or paired with a WT or ectodomain-deleted receptor (Fig. 1E). The membrane integrations of hIL-7R-ΔECD and hγc-ΔECD were confirmed by confocal microscopy (fig. S1D). Among the receptor-expressing cells, the one that co-expressed hIL-7R-ΔECD and hγc-ΔECD exhibited the strongest signaling, reaching ∼40% of ligand-induced STAT5 phosphorylation by the WT receptors (Fig. 1, F and G). This result is consistent with the activation achieved by proteolytic removal of ectodomains (Fig. 1, C and D). Apart from the double ectodomain deletion, the cells expressing hIL-7R-ΔECD or hγc-ΔECD (alone or with a WT receptor) also showed weaker but measurable ligand-independent signaling (Fig. 1, F and G), suggesting that both hIL-7R and hγc ectodomains are involved in receptor autoinhibition in the absence of ligand, i.e., preventing the TMD-ICD regions of hIL-7R and hγc from forming signaling competent complexes. The above results suggest an active role of TMDs in the heterotypic association between hIL-7R and hγc responsible for signaling.

### TMD interactions match the receptor sharing network

We next investigated whether the TMD of the γc (designated γcTMD) can directly associate with that of ILR members (designated IL-xRTMD) in cell membrane and whether the heterodimerization of TMDs also show properties of receptor sharing as the ectodomains. As mentioned above, three shared cytokine receptor families have been identified based on their common chains: γc, βc, and gp130 (*1, 9*). In this study, we focused on the γc family and have selected the TMDs of γc, IL-4R, IL-7R, and IL-9R from both human and mouse for testing oligomerization (Fig. 2A). We also selected the TMD of IL-5R from the βc family to examine for possible non-specific association with γcTMD. The TMD sequences of these receptors (Fig. 2B and fig. S2) do not show any conserved small-xxx-small motif that mediates TMH dimerization in growth factor receptors (*12, 16, 25, 26*). Nor do they show charged residues that mediate intramembrane assembly of activating immunoreceptors (*27-29*).

**Fig. 2.**
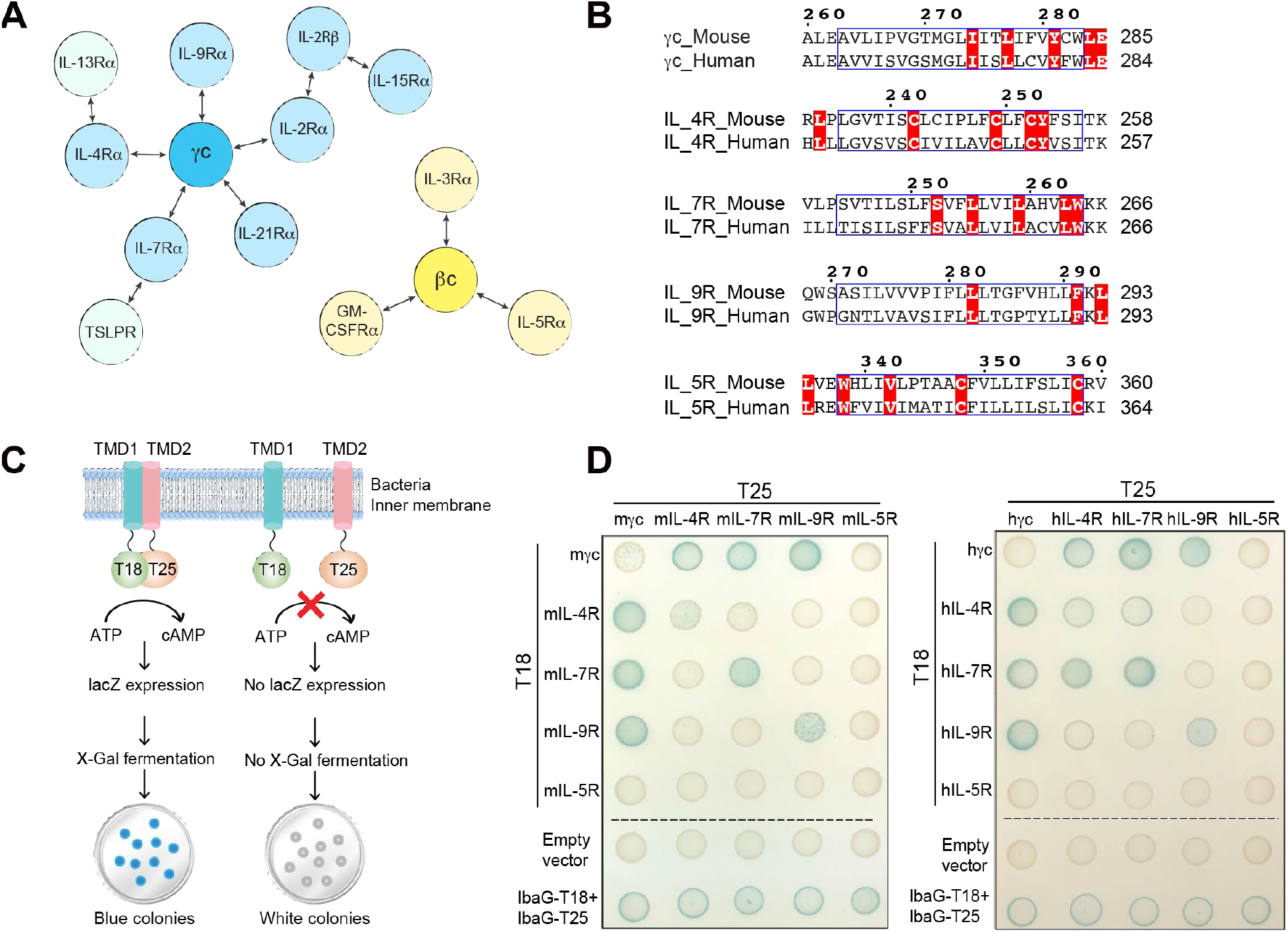
Specific heterodimerization of γcTMD with the TMDs of its family members. (**A**) Schematic diagram showing receptor sharing network for the γ chain (blue) and β chain (yellow) receptor families. Some complexes require a third chain for cytokine recognition and signal transduction, such as IL-2Rα and IL15Rα. IL-4Rα and IL-7Rα can also form heterodimer complexes with IL-13Rα and TSLPR, respectively. (**B**) TMD sequence alignment of representative γc family receptors (γc, IL-4R, IL-7R and IL-9R) and the βc family receptor IL-5R from both mouse and human. Transmembrane regions are highlighted in blue box. 100% conserved residues across species are shown in bold white and red background. (**C**) Schematic illustration of the bacterial two-hybrid (BACTH) system (*30*) for analyzing TMD interaction (see text for details). (**D**) BACTH analysis of TMD interaction for representative γc family receptors and the βc family member IL-5R from both mouse and human. Blue colonies indicate TMD-TMD association in the bacteria inner membrane.

TMD interactions were examined using a bacterial adenylate cyclase two-hybrid (BACTH) assay (*30*). Briefly, two complementary domains (T18 and T25) of the adenylate cyclase (AC) are fused, separately, to the C-termini of two TMDs under investigation and the outer membrane protein A (OmpA) is fused to the TMD N-termini for membrane localization (Fig. 2C). The transmembrane fusion proteins are expressed in a CyaA deficient strain that yields white bacteria colonies by default. Stable association of two TMDs would reconstitute the AC activity, resulting in blue colonies. According to the colony colors, it is obvious that γcTMD can stably heterodimerize with the TMDs of IL-4R, IL-7R, and IL-9R for both mouse and human sequences as the corresponding colonies showed the strongest blue colors (Fig. 2D and fig. S3). In contrast, no association between the TMDs of γc and IL-5R was detected (Fig. 2D and fig. S3), indicating no cross-family TMD association and further validating that TMD association of the γc with its own family members is specific. It is interesting to note that homotypic TMD interaction was also observed for IL-4R, IL-7R, and IL-9R, though on average weaker than the heterotypic interaction involving γcTMD (Fig. 2D). While the heterotypic interactions between γcTMD and IL-xRTMDs are fully consistent with the functionality of receptor sharing in the γc family, potential function of homotypic TMD interactions is unknown.

### Structures of γcTMD in complex with IL-7RTMD and with IL-9RTMD in bicelles

Our observation of specific recognition of the TMDs of IL-4R, IL-7R, and IL-9R by γcTMD was unexpected because their sequences are highly divergent (Fig. 2B and fig. S2). To understand the structural basis of receptor sharing at the membrane level, we determined high resolution NMR structures of two heterodimer complexes, γcTMD bound to IL-7RTMD and γcTMD bound to IL-9RTMD, in DMPC-DHPC bicelles that mimic a lipid bilayer. We first performed an inter-molecular nuclear Overhauser effect (NOE) difference experiment to identify γcTMD residues involved in the heterotypic interactions. In this experiment, (^15^N, ^2^H)-labeled γcTMD was mixed with (^13^C)-labeled IL-7RTMD or IL-9RTMD at 1:1 molar ratio (fig. S4) and ^13^C-attached aliphatic protons of IL-7/9RTMD were used to generate inter-molecular NOEs to ^15^N-attached amide protons of γcTMD (Fig. 3A). The experiment immediately revealed γcTMD residues in the interaction interface. In particular, I274, L277, and I278 experienced inter-molecular NOE both in complex with IL-7RTMD and with IL-9RTMD (Fig. 3A). Full scale structure determination involved measurement of reciprocal inter-chain NOEs between (^15^N, ^2^H)-labeled γcTMD and (^13^C)-labeled IL-7/9RTMD and between (^15^N, ^2^H)-labeled IL-7/9RTMD and (^13^C)-labeled γcTMD (fig. S5 and fig. S6). The final ensemble of structures that satisfy all NMR restraints converged to root mean square deviation (RMSD) of 0.645Å (backbone) and 1.234Å (heavy atoms) for the γcTMD/IL-7RTMD dimer and RMSD of 0.711Å (backbone) and 1.281Å (heavy atoms) for the γcTMD/IL-9RTMD dimer (table S1).

**Fig. 3.**
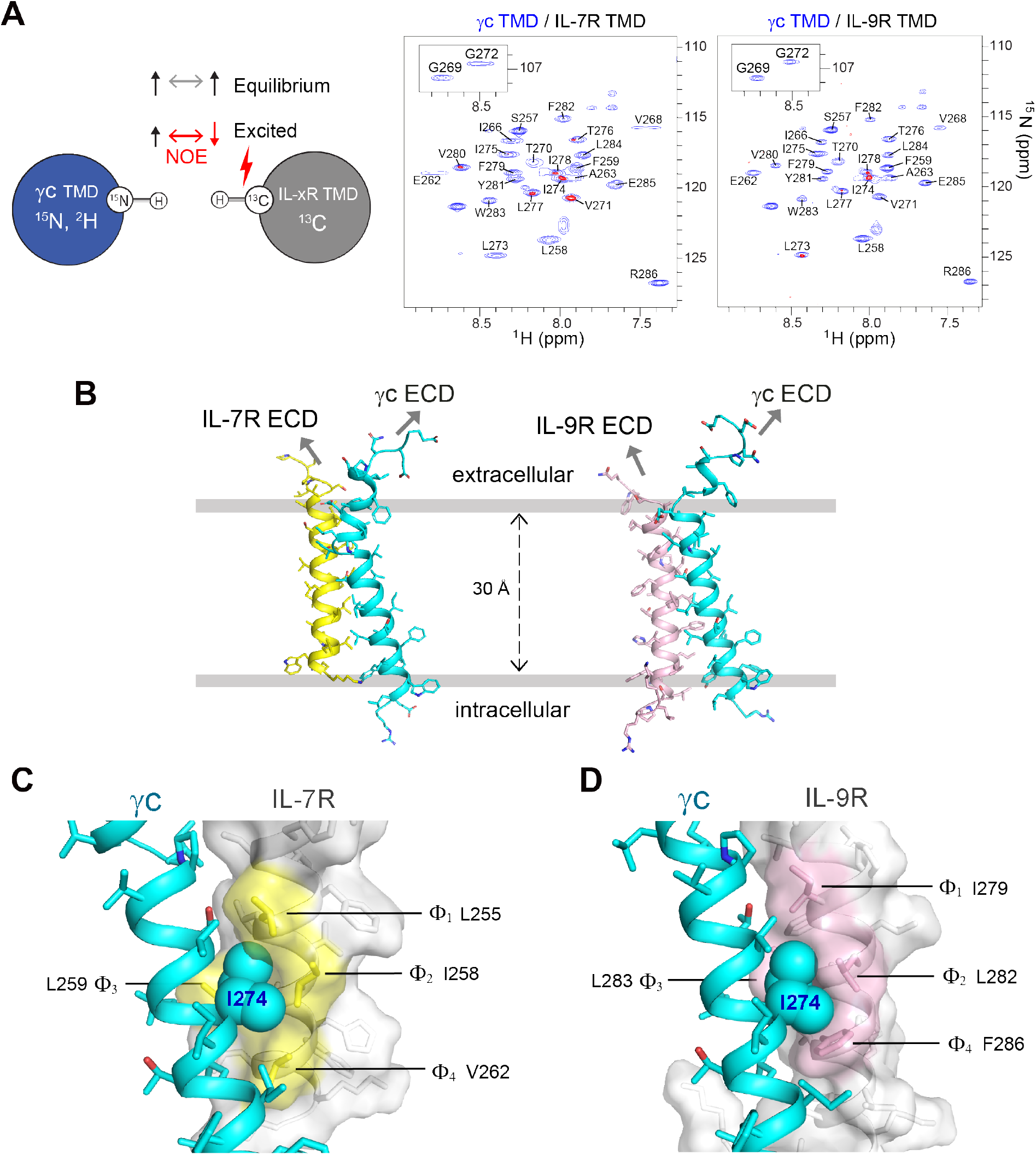
NMR structures of γcTMD in complex with IL-7RTMD and with IL-9RTMD show a common knob-into-hole mechanism of recognition. (**A**) *Left panel*: Detecting residue-specific inter-chain NOEs using the sample of (^15^N, ^2^H) γcTMD mixed with (^13^C) IL-7RTMD or IL-9RTMD at 1:1 molar ratio. The experiment involves recording two interleaved ^1^H-^15^N TROSY-HSQC spectra: one at equilibrium and the other with the ^13^C-attached aliphatic protons inverted during 200 ms of NOE mixing. *Right panel*: Overlaying the difference between two interleaved ^1^H-^15^N TROSY-HSQC spectra (red) onto the reference spectrum (blue) reveals γcTMD residues in close contact with IL-7/9RTMD (right). The spectra were recorded at 303 K and ^1^H frequency of 900 MHz. (**B**) Ribbon representation of the structures of γcTMD in complex with IL-7RTMD (left) and with IL-9RTMD (right) in DMPC-DH_6_PC bicelles with *q* = 0.4. (**C**) A close-up view of γcTMD I274 (sphere) fitting into the hydrophobic pocket of IL-7RTMD (yellow) formed by the Φ_1_-xx-Φ_2_Φ_3_-xx-Φ motif. (**D**) The same close-up view as in (C) of γcTMD I274 fitting into the pocket of IL-9RTMD (pink).

The two transmembrane heterodimer structures are quite homologous despite low sequence identity between IL-7RTMD and IL-9RTMD, and this structural similarity is characterized by ∼20º helical packing angle and the involvement of the same face of γcTMD in packing against IL-7/9RTMD (Fig. 3B). Closer inspection of the two structures identified I274 of γcTMD as the key residue that fills a hydrophobic hole of IL-7/9RTMD comprising four residues (Fig. 3, C and D), reminiscent of the knob-into-hole mechanism. These four residues have the sequence arrangement of Φ_1_-xx-Φ_2_Φ_3_-xx-Φ_4_, where Φ_i_ represents hydrophobic residues such as isoleucine, leucine, valine, and phenylalanine that constitute the pocket. Other γcTMD residues, P267, T270, L277 and Y281, also make close van der Waals (VDW) contact with receptor TMDs but do not appear to be locked in a hole (Fig. 4, A and B). On the side of receptor TMD, S252, L255, L259, and V262 of IL-7RTMD and I279, L282, L283 and F286 of IL-9RTMD are in close VDW contact with γcTMD but none of them are wrapped by a defined pocket (Fig. 4, A and B).

**Fig. 4.**
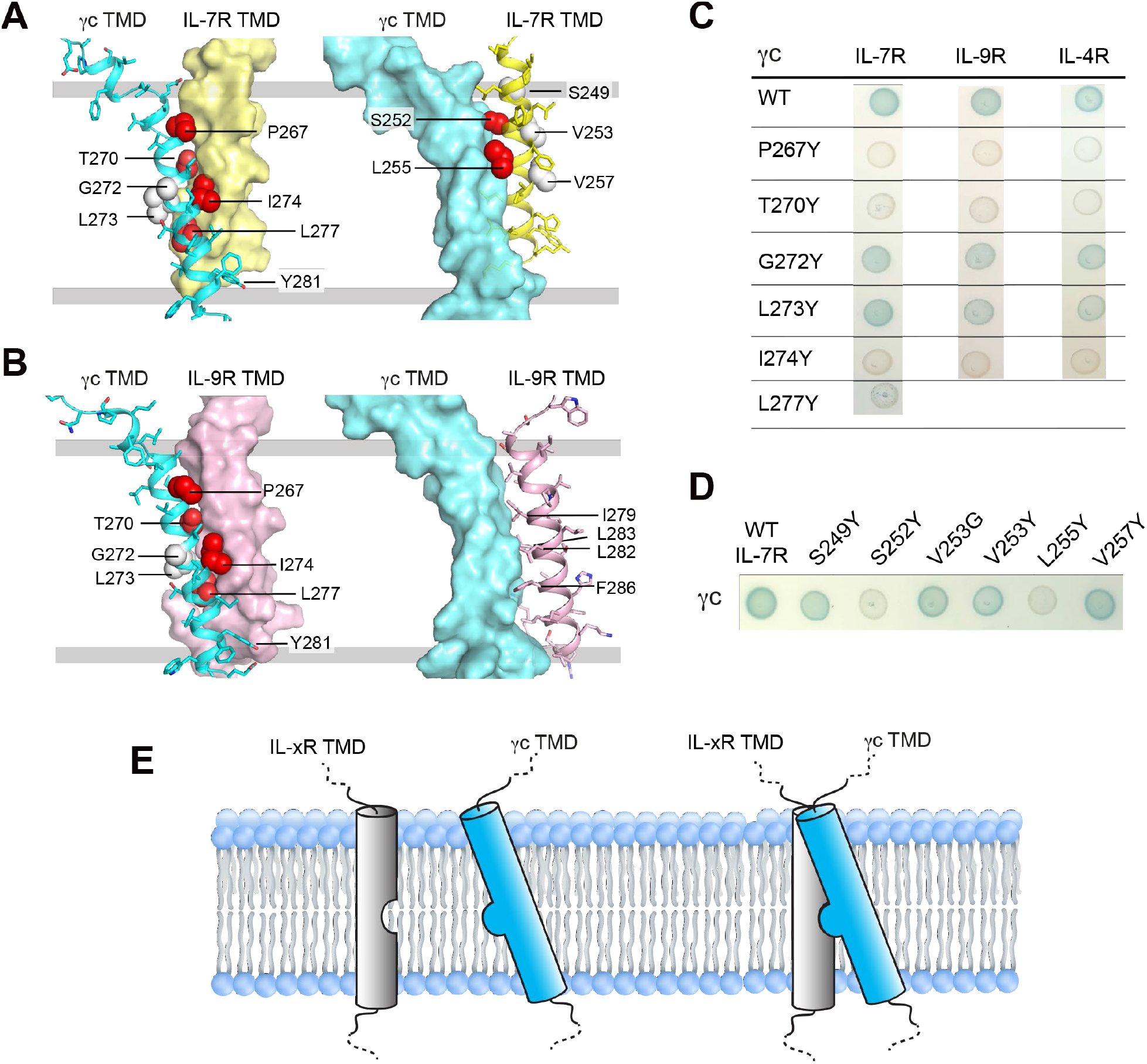
Validation of the surface complementarity and the common mechanism of γcTMD sharing by mutagenesis. (**A**) The contours of the γcTMD / IL-7RTMD heterodimer interface shown with the IL-7RTMD strand surface rendered (left) and the γcTMD surface rendered (right). The γcTMD and IL-7RTMD residues in sphere are residues tested by mutagenesis in (C), with red and white indicating dimer disruption and no effect, respectively. (**B**) Same as in (A) for the γcTMD / IL-9RTMD heterodimer. (**C**) BACTH analysis of the effect of γcTMD mutations on γcTMD association with IL-7RTMD, IL-9RTMD, or IL-4RTMD. Corresponding mutations for human sequences are shown in fig. S7. (**D**) BACTH analysis of the effect of IL-7RTMD mutations on association with γcTMD. (**E**) Schematic illustration of the knob-into-hole mechanism of recognition that mediates receptor sharing within the membrane.

We have examined the key residues in the dimer interface using the BACTH assay and found that mutating P267, T270, I274, or L277 of γcTMD to tyrosine can disrupt heterodimerization with IL-7RTMD (Fig. 4C and fig. S7, A to D) and the same pattern of mutational effect was observed for IL-9RTMD (Fig. 4C and fig. S7, E and F) and IL-4RTMD (Fig. 4C and fig. S7, G and H). On the receptor side, mutating S252 or L255 of IL-7RTMD to tyrosine also abolished heterodimerization (Fig. 4D and fig. S7I). Collectively, the mutagenesis data indicate that the NMR structures are consistent with those expressed in cell membrane while suggesting that intramembrane recognition of the γc by different family members are based largely on the same knob-into-hole mechanism (Fig. 4E).

Another interesting feature of the heterodimer structures is the relatively small binding area, 501.4 Å^2^ between γcTMD and IL-7RTMD or 479.0 Å^2^ between γcTMD and IL-9RTMD, which is consistent with the small number of residues involved in the knob-into-hole interaction. Confinement of interaction to a small area could explain why IL-7RTMD and IL-9RTMD have only 35% identity but both can be recognized by γcTMD. In this regard, the interaction interface between the γc ectodomain and cytokines are also relatively small at ∼1000Å^2^, and the same argument has been proposed for explaining degenerate cytokine recognition by the γc ectodomain (*10*).

### Essential role of the knob-into-hole interaction in membrane for IL-7R signaling

To further examine whether the specific knob-into-hole mechanism of TMD heterodimerization is relevant to receptor signaling, we performed structure guided mutagenesis and tested the mutants using functional assays similar to that used above for receptor activation by TEV protease (Fig. 1, C and D). Based on the structure of γcTMD in complex with IL-7RTMD, the knob residue of γcTMD is I274 in mouse and I273 in human and a key constituent of the hole in IL-7RTMD is L255 (Fig. 5A). These residues were mutated to tyrosine to disrupt the knob-into-hole interaction. In addition, the glycine on the opposite side of the γc TMH (G272 in mouse and G271 in human) and V253 on the opposite side of the IL-7R TMH were selected as no-effect mutations.

**Fig. 5.**
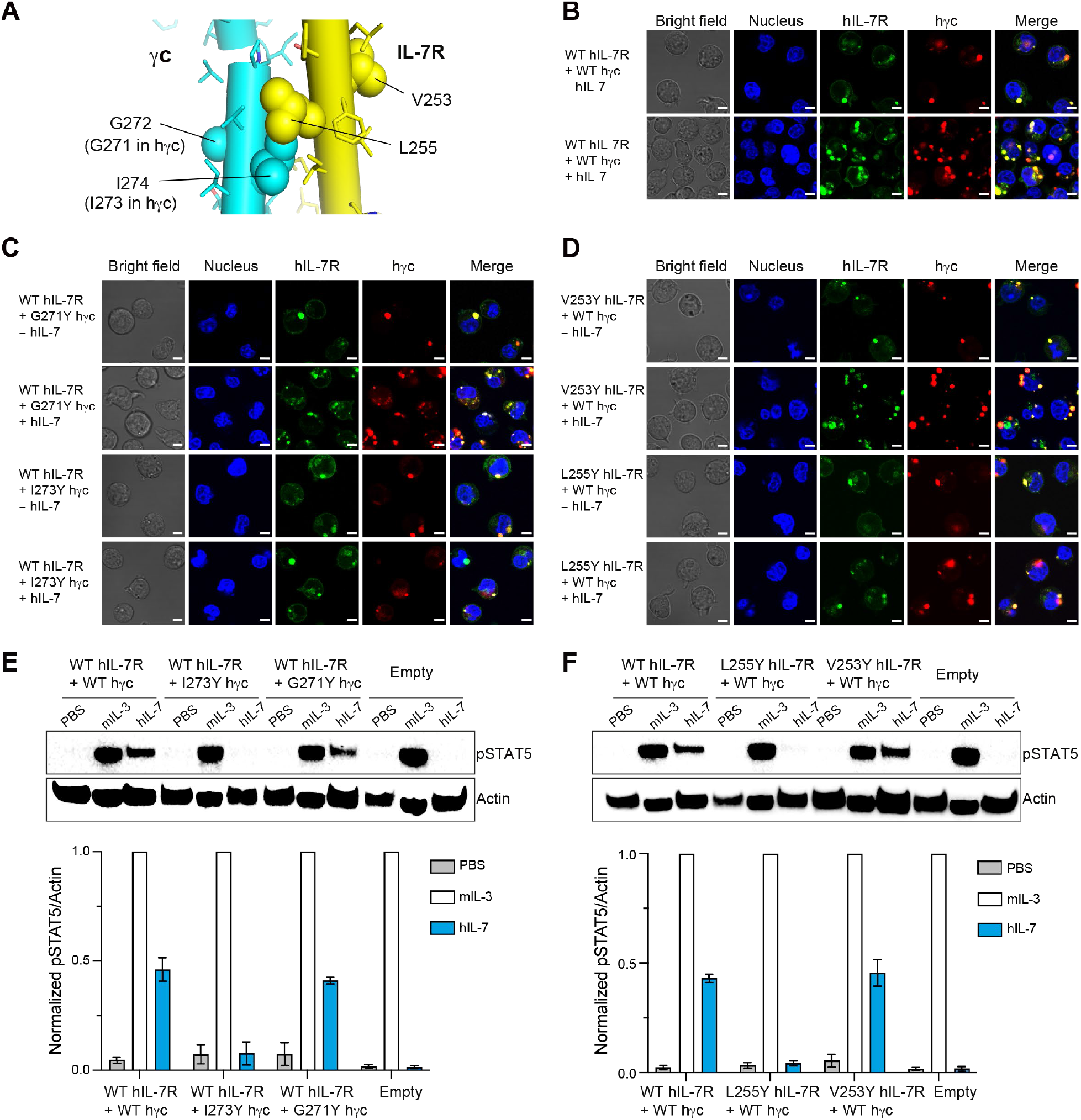
Specific TMD heterodimerization is required for ligand-induced IL-7R signaling. (**A**) Cylinder representation of the γcTMD/IL-7RTMD heterodimer structure showing the positions of the TMD residues of γc and IL-7R (sphere) that were tested for ligand-induced receptor signaling. Specifically, I274 of γcTMD and L255 of IL-7R are important for the knob-into-hole mechanism whereas G272 of γcTMD and V253 of IL-7R are not involved in heterodimerization. (**B**) Co-expression and distribution of WT hIL-7R and hγc on the surface of BaF3 without and with hIL-7 treatment. Cells were stained with DAPI for 1 hour for nucleus staining before imaging. Images were taken with the Olympus Fluoview FV1000 confocal microscope. Scale bar, 5μm. (**C**) Co-expression and distribution of WT hIL-7R and hγc mutants on the surface of BaF3 without and with hIL-7 treatment. Scale bar, 5μm. (**D**) Co-expression and distribution of WT hγc and hIL-7R mutants on the surface of BaF3 without and with hIL-7 treatment. Scale bar, 5μm. (**E**) Phosphorylation of STAT5 in BaF3 cells co-expressing WT hIL-7R with WT hγc or hγc mutants after 5 hours of cytokine deprivation and treatment with PBS, mouse IL-3 (mIL-3) or human IL-7 (hIL-7). Untransfected (empty) BaF3 cells were used as negative control. Signals are detected by immunoblotting (one representative result is shown in upper panel), quantified with pSTAT5/Actin signal ratio (lower panel) and further normalized relative to that induced by mIL-3. Results are from 3 independent experiments (n = 3) and expressed as mean ± SEM. (**F**) Same as in (E) but for BaF3 cells co-expressing WT hγc with WT hIL-7R or hIL-7R mutants.

Full-length hγc and hIL-7R as well as their single-mutants were expressed in BaF3 cells and with a C-terminal fluorescent tag (mCherry for hγc and EGFP for hIL-7R) for monitoring subcellular location in BaF3 cells. Both WT receptors and mutants expressed well on the cell surface and sparse puncta could be observed before ligand addition, suggesting that introduction of the TMD mutations did not affect preligand association (Fig. 5, B to D). Much more puncta appeared after ligand addition for the cells co-expressing WT hIL-7R and G271Y hγc and the cells co-expressing V253Y hIL-7R and WT hγc (Fig. 5, C and D), consistent with IL-7 induced receptor activation. In contrast, the cells expressing either I273Y hγc or L255Y hIL-7R showed no detectable increase of puncta after ligand addition, indicating that breaking TMD heterodimerization strongly attenuated ligand-induced receptor clustering. The imaging results were independently confirmed by immunoblot analysis of STAT5 phosphorylation. As shown in Fig. 5, E and F, disrupting either the knob with I273Y mutation or the hole with L255Y mutation totally abolished signaling whereas G271Y in hγc or V253Y in hIL-7R did not show detectable differences in signaling from the WT receptors. We conclude from the above data that heterotypic association of receptor TMDs mediated by the knob-into-hole mechanism is essential for ligand-induced signaling of IL-7R and this is likely to be applicable to other members of the γc family.

## DISCUSSION

We have shown that the TMDs of IL-7R and IL-9R in the common γc family of cytokine receptors both contain a defined hydrophobic pocket recognized by γcTMD, and this heterotypic recognition within the cell membrane is required for ligand-induced signaling. Previous clinical studies have recorded mutations or frameshifts in the C-terminal half of γcTMD in patients with X-SCID (*21*) that remain incomprehensible because they should not affect cytokine binding or kinase activity.

We find these mutations could be explained by the requirement of TMD heterodimerization in signaling, as the C-terminal half of γcTMD encompasses residues critical for intramembrane pairing of the γc with ILRs (Fig. 3 and Fig. 4). For example, even a single mutation in γcTMD, I274Y, can completely block activation of IL-7R by IL-7 (Fig. 5).

The simple γc TMH is capable of specific recognition of multiple ILR TMHs that have very little sequence homology (Fig. 2 and fig. S2). An intriguing structural question is how specificity and promiscuity are concomitantly achieved. The complementary surface representations of IL-7/9RTMD and γcTMD (Fig. 3) show that whereas IL-7RTMD and IL-9RTMD both have a deep pocket fitting the I274 knob of γcTMD, no defined pockets were observed in γcTMD that fit any of the IL-7/9RTMD residues. In this context, the I274 knob of γcTMD can be perceived as the key that fits the small hydrophobic hole (or lock) presented by the different ILR TMDs, and this could explain the promiscuity. The structural features governing specificity, however, may include beyond the knob-into-hole module. For example, the vertical position in lipid bilayer of the two TMHs are important for aligning the I274 knob with the hydrophobic hole for binding. Other supporting interactions could further enhance specificity, e.g., interaction of L277 and Y281 of γcTMD with complementary surface of the receptor TMH near its C-terminal end (Fig. 4, A and B). Of note, the three residues of γcTMD with 100% conservation across species are I274, L277, and Y281 (Fig. 2B), and these are precisely the residues making close VDW contacts with the receptor TMHs in our structures. The specificity is also manifested by our observation that γcTMD sharing is family specific as γcTMD showed no interaction with the TMD of IL-5R from the βc family (Fig. 2D).

The fact that removal of receptor ectodomains can result in ∼40% of ligand-induced signaling strongly indicates that they are arranged in an autoinhibitory state in the absence of ligand. This autoinhibition likely involves homotypic and heterotypic associations of the ectodomains of ILRs and γc, which is consistent with the observation that co-expression of IL-7R and γc resulted in their co-localization on the cell surface (fig. S1, A and C). The preligand association of ILRs and γc serves autoinhibitory function but could also gather the receptors for efficient ligand engagement. To date, no structural information is available for heterotypic preligand association in the γc receptor family. A crystal structure of homotypic association, however, has been reported for the IL-7R ectodomain (*31*), which shows a head-to-head antiparallel configuration proposed to be autoinhibitory, as it keeps the TMD-ICDs apart (*24*). Of note, our BACTH assay showed that the IL-7RTMD can also homodimerize in addition to heterodimerizing with γcTMD (Fig. 2D). In principle, TMD homodimerization would position the JAK1s closer to favor the “open” state of the kinase for dimerization and activation (*32*), though there has been no report of physiological function of signaling via IL-7R homotypic association other than the gain-of-function disease phenotypes (*33*). It is also unclear whether the TMD homodimerization competes or synergizes with the heterotypic TMD interaction in receptor activation.

Finally, the requirement of specific TMD pairing in signaling of receptors in the common γc family adds another layer of cooperativity to cytokine mediated receptor pairing; it also affords the opportunity to design TMHs to modulate receptor activity. Although protein design of transmembrane protein remains a challenging task, several studies have demonstrated the feasibility of *de novo* design of TMH oligomerization (*34-36*). Indeed, the use of designed TMH to modulate immunoreceptor activity has been explored. For example, transmembrane peptide designed to compete with the TMD heterodimerization between the integrin α and β subunits could push the equilibrium from the inactive to active state (*37*). More recently, designed TMH oligomerization has been applied to mediate chimeric antigen receptor (CAR) oligomerization to enhance the therapeutic window of CAR-T (*38*). The distinct features of the knob-into-hole mechanism that mediates γc sharing and the structural differences among the ILR members of the family could be exploited for designing decoy γcTMDs for selective interference of cytokine signaling.

## Acknowledgement

We thank Qingshan Fu and Wen Chen for help with protein biochemistry, Abdelrahim Zoued and Roney Ian for sharing the materials for BACTH assay, and Jinqian Li for technical support of virus packaging. We thank Can Xie for insightful discussion.

## Author contributions

T.C., R.L.C., H.W., and J.J.C. conceived the study; T.C. and R.L.C. prepared samples and performed NMR analyses; T.C. performed the BACTH and functional cell assays; X.P. helped with cell assays. J.J.C., T.C. and R.L.C. wrote the paper and others helped with editing the paper.

## Data and materials availability

All data are available in the main text or the supplementary materials. The atomic structure coordinate and structural constraints have been deposited in the Protein Data Bank (PDB), accession numbers: 8DDC (γcTMD / IL-7RTMD) and 8DDD (γcTMD / IL-9RTMD). The chemical shift values have been deposited in the Biological Magnetic Resonance Data Bank (BMRB), accession numbers 31026 (γcTMD / IL-7RTMD) and 31027 (γcTMD / IL-9RTMD).

## Supplementary Materials

Materials and Methods

Figs. S1 to S7

Tables S1 to S2

