## Supplemental Information for "Structural basis of *γ*-chain family receptor sharing at the membrane level"

**This PDF file includes:**

Materials and Methods
Figs. S1 to S7
Tables S1 and S2

### Materials and Methods

#### *Cell lines*

*BaF3 cell line*

Cytokine–dependent BaF3 cells were cultured in RPMI 1640 (GIBCO, ThermoFisher Scientific) medium supplemented with 10% fetal bovine serum (GIBCO, ThermoFisher Scientific), 10 ng/mL mouse interleukin-3 (mIL-3) (Sigma) and 100U/ml Pen-Strep (GIBCO, ThermoFisher Scientific), at 37°C, 5% $CO_2$.

*HEK293T cell line*

Human kidney epithelial cell line HEK293T cells (ATCC# CRL-3216; RRID: CVCL_0063) were purchased from ATCC. HEK293T cells were maintained in DMEM (GIBCO, ThermoFisher Scientific) supplemented with 10% fetal bovine serum (GIBCO, ThermoFisher Scientific) and 100U/ml Pen-Strep (GIBCO, ThermoFisher Scientific), at 37°C, 5% $CO_2$.

#### *Bacterial two-hybrid (BACTH) expression and interaction assay*

A bacterial two-hybrid (BACTH) system based on the activity of reconstitution of adenylate cyclase (AC) (*30*) was adopted to address interaction between transmembrane domains (TMDs) in cell membrane. The BACTH kit (Euromedex, EUK001) was originally purchased from Euromedex and kindly provided by Dr. Abdelrahim Zoued and Roney Ian at Harvard Medical School. To detect TMD interaction, two complementary domains (T18 and T25) of the adenylate cyclase (AC) were fused, separately, to the C-termini of two TMDs under investigation and the outer membrane protein A (OmpA) was fused to the TMD N-termini for membrane localization. Two plasmids (OmpA-TMD1-T18, with ampicillin resistance; OmpA-TMD2-T25, with kanamycin resistance) encoding the T18 and T25 fusion proteins were transformed into the CyaA deficient *E.coli* strain BTH101. The co-transformed colonies were selected on LB agar plates containing 50 µg/ml kanamycin and 100 µg/ml ampicillin after incubation at 30°C for 48h. Three colonies for each transformation were picked and cultured in 3ml of LB medium supplemented with 100 µg/ml ampicillin, 50 µg/ml kanamycin and 0.5mM IPTG. After overnight growth of the cultures at 30°C with shaking at 220rpm, 2 µL from each culture were dropped on the reporter LB-

X-GAL plates containing 40 μg/mL X-Gal, 100 μg/ml ampicillin, 50 μg/ml kanamycin and 0.5mM IPTG. The plates were incubated at 30°C for 24h. BolA-like protein IbaG fused with T18 and T25 (IbaG-T18/IbaG-T25), as previously published, was used as a positive control (*39*). Co-expression of the empty vector T18 with T25 was used as a negative control.

***Retroviral production and infection of BaF3 cells***

Wild-type (WT) or mutants of full-length human IL-7R (hIL-7R) were cloned into retroviral vector MSCV-EGFP with the fusion expression of EGFP at the C-terminus. WT or mutants of full-length human γc (hγc) were cloned into retroviral vector MSCV-mCherry with the fusion expression of mCherry at the C-terminus. Plasmids with single site mutations, V253Y and L255Y of hIL-7R and G271Y and I273Y of hγc, were generated by site-directed mutagenesis. All constructs were confirmed by DNA sequencing. Retroviral production and transduction were performed as described previously (*40*). Briefly, HEK293T cells were seeded in 10 cm dishes and cultured until well attached and 60%–70% confluent. 10 μg of retroviral expression plasmid (MSCV- hIL-7R -EGFP or MSCV- hγc-mCherry) and 8 μg of packaging helper plasmid pCL-Eco (Addgene, #12371, Cambridge MA) were co-transfected with Lipofectamine 3000 (ThermoFisher Scientific), following the manufacturer's instructions. 12h after transfection, the medium was replaced with fresh DMEM supplemented with 10% FBS. 24h after changing the medium (36h post transfection), the virus containing supernatant was centrifuged at 1,000g for 10 min at 4°C to remove cells and debris and filtered through a 0.45μm sterile filter. The retrovirus was then concentrated with Retro-X Concentrator according to the manufacturer's instructions.

Next, 100 μl of concentrated retroviral supernatant supplemented with 8 μg/mL polybrene and 10 ng/ml mIL-3 was used to infect 250,000 BaF3 cells. After overnight incubation at 37°C, the medium was replaced with fresh RPMI1640 supplemented with 10% FBS, 10 ng/ml mIL-3 and penicillin-streptomycin. The infected BaF3 cells were cultured until they reached a concentration of $2x10^6$ cells/ml in 10 ml medium. To generate the stable cell line, the co-expressing hIL-7R and hγc cells or single receptor expressing cells were sorted by FACS (Sony SH800Z Cell Sorter) according to the receptor fused EGFP and/or mCherry tags.

***Cytokine-induced activation***

Untransfected (Empty) BaF3 cells and BaF3 stable cell lines co-expressing WT receptors (WT hIL-7R and hγc) or receptors with TEV protease site (TEV-hIL-7R and TEV-hγc) were washed 3 times with PBS followed by cytokine withdraw by culturing cells without cytokine for 5h. After starving, the cells were stimulated with 50 ng/ml mIL-3 or 50 ng/ml hIL-7 for 2h at 37°C and then harvested for immunoblot detection.

***Activation by proteolytic removal of ectodomain***

For testing activation of the IL-7R pathway by proteolytic removal of the ectodomains (ECDs) of IL-7R and γc without ligand, TEV protease cleavage sequence (ENLYFQGGGGGS) was introduced between the ECD and TMD for both hIL-7R (TEV-hIL-7R) and hgc (TEV-hgc). Specifically, the insertion was between residues 235 and 236 of hIL-7R (isoform 1) and between residues 251 and 252 of hγc (isoform 1). BaF3 stable cell lines co-expressing TEV-hIL-7R and TEV-hγc or co-expressing WT hIL-7R and hγc were generated with retroviral infection as described above. The cells were washed 3 times with PBS and cultured without cytokine for 5h. After cytokine withdrawal, untransfected (empty) BaF3 cells, BaF3 cells expressing WT receptors (hIL-7R and hγc) or receptors cleavable by TEV protease (TEV-hIL-7R and TEV-hγc) were treated with 0, 10, 20, 30, 40, 50, 60, 100 or 200 μg/ml TEV enzyme overnight at 37°C. The cells were then harvested and the activation of the IL-7R signaling pathway was evaluated by immunoblotting of phosphorylation of the STAT5. Results were from 3 independent experiments (n = 3), and expressed as mean ± SEM.

***Activation due to ectodomain deletion***

The hIL-7R with ECD (residues 24 to 235) deleted (hIL-7R-ΔECD) and the hγc with ECD (residues 26 to 251) deleted (hγc-ΔECD) were subcloned into MSCV-EGFP and MSCV-mCherry retroviral vectors, respectively. Retroviral production and infection were performed as described above. BaF3 stable cell lines expressing hIL-7R-ΔECD or hγc-ΔECD (alone or with a WT receptor) were washed 3 times with PBS followed by cytokine withdrawal by culturing cells without cytokine overnight. The cells were then harvested for immunoblot detection.

***Cell imaging***

BaF3 stable cell lines expressing the WT, mutant, or ECD-deleted hIL-7R and hγc receptors were washed 3 times with PBS followed by cytokine withdrawal by culturing cells without cytokine for 5h. Observation of receptors without and with cytokine stimulation was undertaken by stimulating cells with either PBS or 50 ng/ml hIL-7 for 2h. For observation of receptor activation by TEV protease, cells were incubated with either PBS, PBS with 50 ng/ml human hIL-7 or PBS with 50 μg/ml TEV enzyme overnight at 37°C. All the cells were transferred to glass-bottomed dishes coated with poly-glycine and stained with 5 μM DAPI for 1h before imaging. All the images were taken with Olympus Fluoview FV1000 confocal microscope.

***Immunoblotting***

Western blotting (WB) was used to detect the phosphorylation of STAT5 as described previously (*33*). Briefly, the harvested cells were washed 2 times with ice-cold PBS and lysed for 20min in ice-cold RIPA lysis and extraction buffers supplemented with Halt protease inhibitor cocktail and Halt phosphatase inhibitor cocktail (ThermoFisher Scientific), respectively. The lysates were centrifuged at 14,000rpm for 20min at 4°C. The supernatants were transferred to clean tubes and NuPAGE LDS Sample Buffer (4X) (ThermoFisher Scientific) was added. After incubation at 95°C for 5 minutes, the supernatants were subject to 12% SDS-PAGE (Genscript) and transferred to PVDF membranes (Thermo Scientific). The PVDF membranes were then blocked with 5% BSA in TBS-Tween buffer (20 mM Tris, 150 mM NaCl, 0.1% Tween 20, pH 7.6) overnight at 4°C, followed by incubation with primary antibodies against Phospho-STAT5 (Tyr694) (C11C5) (Cell Signaling Technology, #9359) and Beta-actin (Cell signaling, #4970S) with 1:1000 dilution for 1h at room temperature. After washing with TBST buffer 3 times, the membranes were incubated with horseradish-peroxidase–conjugated secondary antibody (Cell signaling, #7074S) for 40min at room temperature. The membranes were washed with TBST buffer again 6 times and the WB signal was detected with Pierce ECL Western blotting substrate (ThermoFisher Scientific) following the product's instruction.

***Quantification and statistical analysis***

The intensity of WB bands was measured with ImageJ for quantification. Statistical significance was calculated with GraphPad Prism 9.3.1 (unpaired Student's test) and shown in each figure or figure legend.

***Transmembrane domain (TMD) sample preparation for NMR investigations***

TMD expression, purification, and reconstitution in bicelles were undertaken following a previously published protocol (*41*).

*Expression of fusion protein*

The DNA sequences coding for the TMDs of mouse γc (residues 253-286), IL-7R (residues 236-266) and IL-9R (residues 268-298) were synthesized by GenScript, codon optimized for E.*coli* expression. For the γc, amino acids M271 and C282 were mutated to valine and phenylalanine, respectively, based on sequence alignment across species for purification and better expression. Since TMDs used for NMR studies were all from mouse, γcTMD, IL-7RTMD and IL-9RTMD refer to sequences from mouse unless otherwise indicated. The DNAs were cloned into the pMM-LR6 vector, fused at the C-terminus of the $His_9$-TrpLE tag sequence with a methionine codon between the tag and the TMD sequence for tag cleavage. The TrpLE tag enables TMDs, which are very hydrophobic and potentially toxic to cells, to be driven into inclusion bodies, resulting in higher expression yields. BL21 DE3 (New England Biolabs) cells were transformed with the plasmids containing gcTMD or IL-7RTMD gene sequences following manufacturer's recommendations, whereas BL21 C43 cells were transformed with the IL-9RTMD containing plasmid and selected using 50 μg/mL kanamycin. Precultures were grown in 100mL of M9 minimal medium. Depending on the isotopic labelling required, M9 medium was composed of $H_2O$, 50mM $Na_2HPO_4$, 25mM $KH_2PO_4$, 10mM NaCl, 2mM $MgSO_4$, 0.1 mM $CaCl_2$, 4g/L D-Glucose (Cambridge Isotope Laboratories, ref CLM-13965 for $^{13}C$ D-glucose), 1g/L of ammonium chloride (Cambridge Isotope laboratories, ref NLM-467-5 for $^{15}N$ ammonium chloride) and 50 μg/mL kanamycin sulphate. For perdeuterated sample (e.g., ($^{15}N$, $^{2}H$)-labeled sample), the M9 medium comprised 99.8% $D_2O$ (Cambridge Isotope Laboratories DLM-4-99.8-1000) and 4g/L deuterated glucose (D1,2,3,4,5,6,7 D-Glucose, ref DLM-20621) with all other components the same as above. Precultures were incubated at 37°C and 220rpm, then used to inoculate 900mL of the same medium and incubated further until the $OD_{600}$ reached between 0.6 and 0.8. Protein expression was induced with 0.5mM isopropyl β-D-thiogalatopyranoside (IPTG, Sigma-Aldrich) for IL-7RTMD and γcTMD or 1mM IPTG for IL-9RTMD. Expression was undertaken at 37°C overnight for mIL-7RTMD and IL-9RTMD, and 22°C overnight for γcTMD (~18h).

*Purification of fusion protein*

Cells were harvested by centrifuging 30min at 4,000rpm and 4°C, and resuspended in lysis buffer (50 mM Tris, pH 8.0, 200 mM NaCl). Cells were then sonicated on ice for 10min at 1 sec intervals, 50% maximum amplitude and the resulting lysate centrifuged 30min at 35,000g and 4°C. The pelleted inclusion bodies were resuspended in guanidine buffer (6M guanidine–HCl, 50mM Tris, pH 8.0, 200mM NaCl, filtered through a 0.22mm MCE membrane, (MF-Millipore™), and 1% (vol/vol) Triton X-100 added) and dissolved using a glass tissue grinder until homogenous. The lysate was centrifuged one last time 20min at 35,000g and 4°C, and the supernatant was added to Ni-NTA resin (HisPur™Ni-NTA resin, Thermo Scientific), thoroughly washed with $dH_2O$ beforehand. The solution was left to incubate on a rotator gently for at least two hours or overnight. The mixture was transferred to a glass chromatography column and purified by gravity flow. The flow through was discarded, and the resin was washed twice with 10 resin volumes of 8M urea, and then washed twice with 10 resin volumes of $dH_2O$. The target protein was eluted in 4 consecutive resin volumes of 90% formic acid.

*Release of TMD by cyanogen bromide cleavage*

Cyanogen Bromide (CNBr) was used to cleave the His9-TrpLE tag from the TMD by hydrolyzing the peptide bond at the C-terminus of the methionine residue. (It is therefore imperative that the TMD target sequence not contain any other methionine). Room temperature cyanogen bromide was added directly to the eluate (0.2 g/mL), and vortexed until completely dissolved. The solution was left at room temperature, shielded from light, and under a gentle nitrogen stream for 1h. The reaction time here was carefully controlled to avoid formylation of the TMD. The mixture was then dialyzed twice against 100x volumes of water, for 40min each time, in a 3.5 kDa dialysis cassette (Thermo Scientific). After dialysis, the solution was added with $dH_2O$ to aid freezing and lyophilized until completely dry (about two full days).

*TMD isolation by HPLC*

Once fully lyophilized into a dry powder, the protein was dissolved with 4mL of 90% formic acid and injected into a HPLC system. Purification was undertaken using a Zorbax 300SB-C3 column, 5mm, 9.4x250mm (Agilent), pre-equilibrated with a buffer of 5% isopropanol (Merck), 95% $dH_2O$ and 0.1% trifluoroacetic acid (Merck). Elution was performed with a gradient of 30 to 100% elution buffer (25% acetonitrile (Merck), 75% isopropanol, 0.1% trifluoroacetic acid) in 140mL at 2 mL/min. Elution peaks corresponding to the TMD of interest were collected and lyophilized. When making a mixed sample with two differently labelled TMDs, they were mixed at an approximative 1:1 molar ratio after elution from Ni-NTA resin by formic acid and before cyanogen bromide cleavage. During the HPLC purification, the TMDs eluted as two separate peaks. The absorbance of the eluate was measured for each peak and they were mixed, again, at a 1:1 molar ratio after measurement of absorbance at 280nm using a Nanodrop and prior to lyophilization.

***Reconstitution of TMD in bicelles***

About 1 to 2mg of dried TMD powder was resuspended in HFIP (1,1,1,3,3,3-Hexafluoro-2-propanol, Oakwood Chemical). Meanwhile, 9mg of DMPC (14:0 PC 1,2-dimyristoyl-sn-glycero-3-phosphocholine, Avanti Polar Lipids) or d54-DMPC (Cortecnet, ref CD5012P025) if using deuterated, and 27mg of DHPC (06:0 PC 1,2-dihexanoyl-sn-glycero-3-phosphocholine, Avanti Polar Lipids) or d26-DHPC if using deuterated (Cortecnet, ref CD5010P025) were measured and dissolved in HFIP until homogenous. If reconstituting the γcTMD and IL-9RTMD together, the amount of lipid and detergent was doubled. The protein mixture was then added to the lipid/detergent mixture and the solution was dried under a gentle nitrogen stream for ~1h. Drying was pursued overnight by lyophilization to remove any organic solvent. The dried film was resuspended in 3mL of 8M urea and dialyzed for a total of 6h against 1L of NMR buffer (50mM phosphate buffer pH 6.7) in a 3.5 kDa dialysis cassette, changing the bath every two hours. DHPC was added regularly to the cassette to replace the detergent lost during dialysis (about 3-5mg lost per hour). The protein was then concentrated in a 3 kDa centrifugal filter (Amicon Ultra 4, Ultracel 3K) at 4000 rpm until the concentration reached about 0.5mM per TMD. $D_2O$ was added at 10% of final volume (for frequency locking). The ratio ($q$) of DMPC to DHPC was verified by integrating the corresponding methyl peaks acquired by 1D $^1H$ NMR spectrum and adjusted accordingly for the $q$ = 0.4-0.5. At this ratio, the diameter of the planar region of the bicelle is 40-45 Å.

#### *Assignment of NMR Resonances*

NMR data were collected at 303K on Bruker spectrometers operating at $^{1}$H frequency of 900MHz, 800MHz, 700MHz, or 600MHz equipped with cryogenic probes. NMR data sets were processed using nmrPipe (*42*). NMR spectra were analyzed using XEASY (*43*) and CcpNmr Analysis v.2 (*44*). For sequence-specific assignment of backbone $^{1}H^{N}$, $^{15}$N, $^{13}C^{\alpha}$ and $^{13}$C' resonances of the three TMDs in this study, ($^{15}$N, $^{13}$C, 85% $^{2}$H)-labeled γcTMD, IL-7RTMD and IL-9RTMD were purified and reconstituted in bicelles separately. Each of the three samples were used to record 3D TROSY-based HNCA, HN(CO)CA, HN(CA)CO and HNCO spectra (*45, 46*) at $^{1}$H frequency of 600MHz. Complete assignment of the structured region was achieved for all three TMDs. Similarly, ($^{15}$N, $^{13}$C)-labeled γcTMD, IL-7RTMD and IL-9RTMD were purified and reconstituted in deuterated bicelles separately for the assignment of aliphatic and aromatic resonances of protein sidechains. Each of the three samples were used to record a 3D $^{15}$N-edited NOESY-TROSY-HSQC (τNOE = 60ms) and a 3D $^{13}$C-edited NOESY-HSQC (τNOE = 100ms) spectra at $^{1}$H frequency of 700MHz. The aliphatic proton resonances were assigned by comparing NOE patterns (specific to helix structure) in $^{15}$N-edited and $^{13}$C-edited NOE strips. Finally, both backbone and sidechain assignments of γcTMD, IL-7RTMD, or IL-9RTMD alone were traced when mixed with its binding partner, which were further confirmed by NOE analysis of the heterodimeric complexes (see below).

#### *Assignment of NOE Restraints*

Due to the complexity associated with heterodimerization, assignment of intermolecular NOEs was challenging as it required preparation of multiple samples with different isotope labeling schemes in deuterated bicelles. For NOE analysis of γcTMD in complex with IL-7RTMD, we prepared samples of ($^{15}$N, $^{2}$H)-labeled γcTMD mixed with ($^{13}$C)-labeled IL-7RTMD at 1:1 ratio and ($^{15}$N, $^{2}$H)-labeled IL-7RTMD mixed with ($^{13}$C)-labeled γcTMD for detecting complementary inter-chain NOEs. We first confirmed TMD heterodimerization in bicelles by performing a 2D NOE difference experiment at 700MHz, which involved recording two interleaved $^{1}$H-$^{15}$N

TROSY-HSQC spectra: one preceded by 200ms of NOE mixing at spin equilibrium (magnetization of amide and aliphatic protons in the same direction; spectrum A), and the other preceded by 200ms of NOE mixing with the magnetization of $^{13}$C-attached aliphatic protons selectively inverted by 8ms of $^{1}J_{CH}$ modulation (spectrum B). Inter-chain NOEs between $^{15}$N-attached protons of one chain and the $^{13}$C-attached methyl protons of the opposing chain were revealed by subtracting spectrum A from B. Upon confirmation of specific heterotypic interaction, we then performed 3D $^{15}$N-edited NOESY-TROSY (τNOE = 200 ms) and 3D $^{13}$C-edited NOESY-HSQC (τNOE = 200 ms) at $^{1}$H frequency of 900MHz for each of the two samples above to assign the inter-chain NOE peaks. These complementary and reciprocal inter-chain NOEs identified the helix-helix packing interface. In addition to inter-chain NOEs between amide and methyl protons, we recorded another set of 3D $^{15}$N-edited NOESY-TROSY (τNOE = 100 ms) and 3D $^{13}$C-edited NOESY-HSQC (τNOE = 150 ms) at 900MHz of a uniformly labeled sample with ($^{15}$N, $^{13}$C)-labeled γcTMD mixed with ($^{15}$N, $^{13}$C)-labeled IL-7RTMD at 1:1 ratio. This set of NOE data was used to assign local NOEs for the two TMDs as well as inter-chain NOEs between aliphatic protons.

For NOE analysis of γcTMD in complex with IL-9RTMD, we prepared samples of ($^{15}$N, $^{2}$H)-labeled γcTMD mixed with ($^{13}$C)-labeled IL-9RTMD at 1:1 ratio and ($^{15}$N, $^{2}$H)-labeled IL-9RTMD mixed with ($^{13}$C)-labeled γcTMD for detecting complementary inter-chain NOEs. Again, we first confirmed TMD heterodimerization in bicelles by performing the 2D NOE difference experiment described above at 700MHz. After confirming specific heterodimerization, we performed 3D $^{15}$N-edited NOESY-TROSY (τNOE = 200 ms) and 3D $^{13}$C-edited NOESY-HSQC (τNOE = 200 ms) at $^{1}$H frequency of 800MHz for each of the two samples above to assign the inter-chain NOE peaks. These complementary and reciprocal inter-chain NOEs identified the helix-helix packing interface for the γcTMD – IL-9RTMD interaction.

***Structure Calculation***

The structures were generated using the program XPLOR-NIH (*47*). First, the monomer structures of γcTMD, IL-7RTMD, and IL-9RTMD were generated using the backbone dihedral restraints derived from $^{15}$N, $^{1}$H$^{N}$, $^{13}$Cα, and $^{13}$C' chemical shifts using the TALOS+ program (*48*). Second, the monomer structures and inter-chain NOEs between amide and methyl protons were used to generate crude structures of γcTMD/IL-7RTMD and γcTMD/IL-9RTMD heterodimers. Finally, the initial dimer structures were fed to the XPLOR-NIH for iterative refinement against all NMR restraints, while assigning more self-consistent inter- and intra- chain NOEs in both $^{13}C$-edited NOESY-HSQC and isotopically mixed NOE spectra after each iteration. This iterative process resulted in ~5 NOE restraints per residue for the structured regions of the two heterodimer complexes (table S1).

The XPLOR refinement used a simulated annealing (SA) protocol in which the temperature in the bath was cooled from 1000 to 200K with steps of 20K. The NOE restraints were enforced by flat-well harmonic potentials, with the force constant ramped from 2 to 30 kcal/mol $Å^{-2}$ during annealing. Backbone dihedral angle restraints were taken from the 'GOOD' dihedral angles from TALOS+ (*48*), all with a flat-well (± the corresponding uncertainties from TALOS+) harmonic potential with force constant ramped from 5 to 1000 kcal/mol $rad^{-2}$. For each of the two heterodimers, a total of 100 structures were calculated and 15 lowest energy structures were selected as the final structural ensemble (fig. S5D and S6D; table S1).

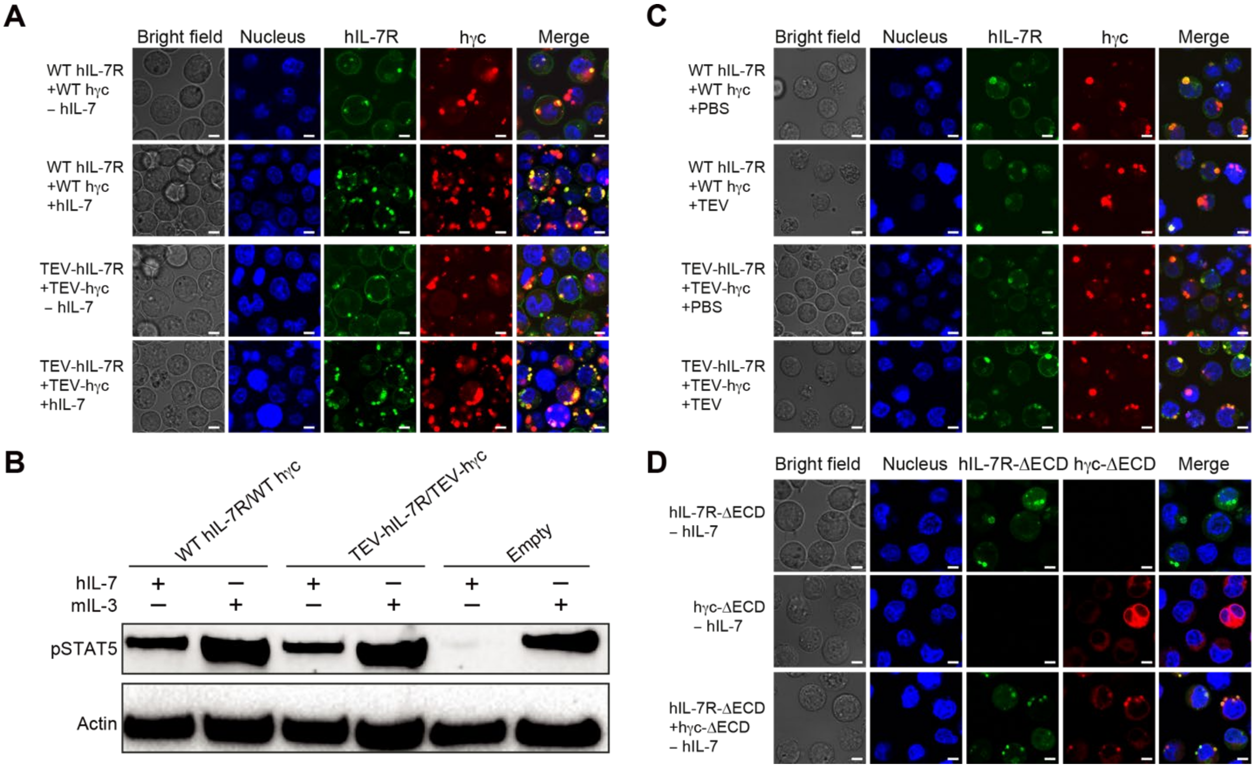


**Fig. S1. Cell surface distribution and activation of IL-7R and γc.**

(**A**) Confocal images of BaF3 cells expressing WT receptors (WT IL-7R / WT γc) or TEV-site-inserted receptors (TEV-IL-7R / TEV-γc) without and with 50 ng/mL hIL-7. Cells were stained with DAPI for 1h for nucleus staining before imaging. Images were taken with the Olympus Fluoview FV1000 confocal microscope. Scale bar, 5μm. (**B**) IL-7R signaling pathway activation detected by STAT5 phosphorylation in untransfected (Empty) BaF3 cells or BaF3 cells expressing WT receptors (WT IL-7R / WT γc) or TEV-site-inserted receptors (TEV-IL-7R / TEV-γc) after treatment with 50 ng/mL hIL-7 or mIL-3. (**C**) Confocal images of BaF3 cells expressing WT receptors (WT IL-7R / WT γc) or TEV-site-inserted receptors (TEV-IL-7R /TEV-γc) without and with 50 μg/mL TEV protease. Scale bar, 5μm. (**D**) Confocal images of BaF3 cells expressing ECD-deleted IL-7R (IL-7R-ΔECD) or ECD-deleted γc (γc-ΔECD) or co-expressing IL-7R-ΔECD with (γc-ΔECD) in the absence of cytokine. Scale bar, 5μm.

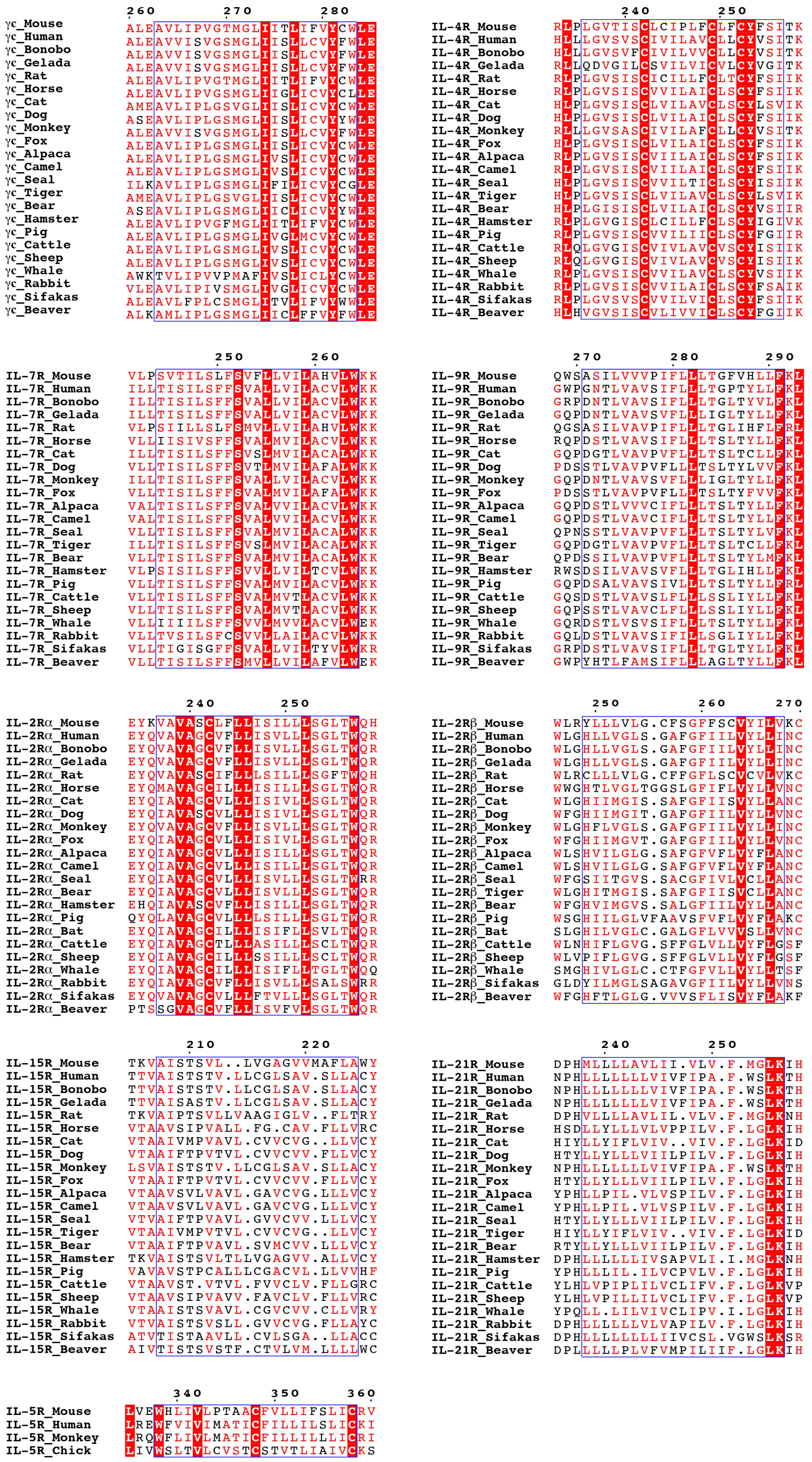
260 270 280
γc_Mouse ALEAVLIPVGTMGLIITLIFVYCWLE
γc_Human ALEAVVISVGSMGLIISLLCVYFWLE
γc_Bonobo ALEAVVISVGSMGLIISLLCVYFWLE
γc_Gelada ALEAVVISVGSMGLIISLLCVYFWLE
γc_Rat ALEAVLIPVGTMGLIITLIFVYCWLE
γc_Horse ALEAVLIPVGSMGLIIGLICVYCLLE
γc_Cat AMEAVLIPLGSVGLIISLICVYCWLE
γc_Dog ASEAVLIPLGSMGLIISLICVYYWLE
γc_Monkey ALEAVVISVGSMGLIISLLCVYFWLE
γc_Fox ALEAVLIPLGSMGLIISLICVYCWLE
γc_Alpaca ALEAVLIPLGSMGLIVSLICVYCWLE
γc_Camel ALEAVLIPLGSMGLIVSLICVYCWLE
γc_Seal ILKAVLIPLGSMGLIFILICVYCGLE
γc_Tiger AMEAVLIPLGSVGLIISLICVYCWLE
γc_Bear ASEAVLIPLGSMGLIICLICVYYWLE
γc_Hamster ALEAVLIPVGFMGLIITLIFVYCWLE
γc_Pig ALEAVLIPLGSMGLIVGLMCVYCWLE
γc_Cattle ALEAVLIPLGSMGLIVSLICVYCWLE
γc_Sheep ALEAVLIPLGSMGLIVSLICVYCWLE
γc_Whale AWKTVLIPVVPMAFIVSLICLYCWLE
γc_Rabbit VLEAVLIPIVSMGLIVGLICVYCWLE
γc_Sifakas ALEAVLFPLCSMGLITVLIFVYWWLE
γc_Beaver ALKAMLIPLGSMGLIICLFFVYFWLE
240 250
IL-4R_Mouse RLPLGVTISCLCIPLFCLFCYFSITK
IL-4R_Human HLLLGVSVSCIVILAVCLLCYVSITK
IL-4R_Bonobo HLLLGVSVFCIVILVVCLLCYVSITK
IL-4R_Gelada RLLQDVGILCSVILVICVLCYVGITK
IL-4R_Rat RLPLGVSISCICILLFCLTCYFSIIK
IL-4R_Horse RLPLGVSISCVVILAICLSCYFSIIK
IL-4R_Cat HLPLGVSISCLVILAVCLSCYLSVIK
IL-4R_Dog HLPLGVSISCLVILAICLSCYFSIIK
IL-4R_Monkey RLLLGVSASCIVILAFCLLCYVSITK
IL-4R_Fox HLPLGVSISCLVILAICLSCYFSIIK
IL-4R_Alpaca RLPLGVSISCVIILAICLSCYFSIIK
IL-4R_Camel RLPLGVSISCVIILAICLSCYFSIIK
IL-4R_Seal HLPLGVSISCVVILTICLSCYISIIK
IL-4R_Tiger HLPLGVSISCLVILAVCLSCYLSVIK
IL-4R_Bear HLPLGISISCLVILAICLSCYVSIIR
IL-4R_Hamster RLPLGVGISCLCILLFCLSCYIGIVK
IL-4R_Pig RLPLGVSISCVVILIICLSCYFGIIR
IL-4R_Cattle RLQLGVGISCVIVLAVCVSCYISIIK
IL-4R_Sheep RLQLGVGISCVIVLAVCVSCYISIIK
IL-4R_Whale RLPLGVSISCVVILAVCLSCYVSIIK
IL-4R_Rabbit RLPLGVSISCVVILAICLSCYFSAIK
IL-4R_Sifakas RLPLGVSVSCVVILVICLSCYFSIIK
IL-4R_Beaver HLHVGVSISCVLIVVICLSCYFGIIK
250 260
IL-7R_Mouse VLPSVTILSLFSVFLLVILAHVLWKK
IL-7R_Human ILLTISILSFFSVALLVILACVLWKK
IL-7R_Bonobo ILLTISILSFFSVALLVILACVLWKK
IL-7R_Gelada ILLTISILSFFSVALLVILACVLWKK
IL-7R_Rat VLPSIILLSLFSMVLLVILAHVLWKK
IL-7R_Horse VLLIISIVSFFSVALMVILACVLWKK
IL-7R_Cat ILLTISILSFFSVSLMVILACALWKK
IL-7R_Dog VLLTISILSFFSVTLMVILAFALWKK
IL-7R_Monkey ILLTISILSFFSVALLVILACVLWKK
IL-7R_Fox VLLTISILSFFSVALMVILAFALWKK
IL-7R_Alpaca VALTISILSFFSVALVVILACVLWKK
IL-7R_Camel VALTISILSFFSVALVVILACVLWKK
IL-7R_Seal VLLTISILSFFSVALMVILACALWKK
IL-7R_Tiger ILLTISILSFFSVSLMVILACALWKK
IL-7R_Bear VLLTISILSFFSVALMVILACALWKK
IL-7R_Hamster VLPSISILSFFSVVLLVILTCVLWKK
IL-7R_Pig VLLTISILSFFSVALMVILACVLWKK
IL-7R_Cattle VLLTISILSFFSVALMVTLACVLWKK
IL-7R_Sheep VLLTISILSFFSVALMVTLACVLWKK
IL-7R_Whale VLLIIIILSFFSVVLMVVLACVLWEK
IL-7R_Rabbit VLLTVSILSFCSVVLLAILACVLWKK
IL-7R_Sifakas VLLTIGISGFFSVALLVILTYVLWKR
IL-7R_Beaver VLLTISILSFFSMVLLVILAFVLWEK
270 280 290
IL-9R_Mouse QWSASILVVVPIFLLLTGFVHLLFKL
IL-9R_Human GWPGNTLVAVSIFLLLTGPTYLLFKL
IL-9R_Bonobo GRPDNTLVAVSIFLLLTGLTYVLFKL
IL-9R_Gelada GQPDNTLVAVSVFLLLIGLTYLLFKL
IL-9R_Rat QGSASILVAVPIFLLLTGLIHFLFRL
IL-9R_Horse RQPDSTLVAVSIFLLLTSLTYLLFKL
IL-9R_Cat GQPDGTLVAVPVFLLLTSLTCLLFKL
IL-9R_Dog PDSSTLVAVPVFLLLTSLTYLVVFKL
IL-9R_Monkey GQPDNTLVAVSVFLLLIGLTYLLFKL
IL-9R_Fox PDSSTLVAVPVFLLLTSLTYFVVFKL
IL-9R_Alpaca GQPDSTLVVVCIFLLLTSLTYLLFKL
IL-9R_Camel GQPDSTLVAVCIFLLLTSLTYLLFKL
IL-9R_Seal QPNSSTLVAVPVFLLLTSLTYLVFKL
IL-9R_Tiger GQPDGTLVAVPVFLLLTSLTCLLFKL
IL-9R_Bear QPDSSILVAVPVFLLLTSLTYLMFKL
IL-9R_Hamster RWSDSILVAVSVFLLLTGLIHLLFKL
IL-9R_Pig GQPDSALVAVSIVLLLTSLTYLLFRL
IL-9R_Cattle GQSDSTLVAVSLFLLLSSLIYLLFKL
IL-9R_Sheep GQPSSTLVAVCLFLLLSSLIYLLFKL
IL-9R_Whale GQRDSTLVSVSIFLLLTSLTYLLFKL
IL-9R_Rabbit GQLDSTLVAVSIFILLSGLIYLLFKL
IL-9R_Sifakas GRPDSTLVAVSIFLLLTGLTYLLFKL
IL-9R_Beaver GWPYHTLFAMSIFLLLAGLTYLLFKL
240 250
IL-2Rα_Mouse EYKVAVASCLFLLISILLLSGLTWQH
IL-2Rα_Human EYQVAVAGCVFLLISVLLLSGLTWQR
IL-2Rα_Bonobo EYQVAVAGCVFLLISVLLLSGLTWQR
IL-2Rα_Gelada EYQVAVAGCVFLLISVLLLSGLTWQR
IL-2Rα_Rat EYQVAVASCIFLLLSILLLSGFTWQH
IL-2Rα_Horse EYQMAVAGCILLLISILLLSGLTWQR
IL-2Rα_Cat EYQIAVAGCVLLLISIVLLSGLTWQR
IL-2Rα_Dog EYQIAVASCVLLLISIVLLSGLTWQR
IL-2Rα_Monkey EYQVAVAGCVFLLISVLLLSGLTWQR
IL-2Rα_Fox EYQIAVAGCVLLLISIVLLSGLTWQR
IL-2Rα_Alpaca EYQIAVAGCVLLLISILLLSGLTWQR
IL-2Rα_Camel EYQIAVAGCVLLLISILLLSGLTWQR
IL-2Rα_Seal EYQIAVAGCVLLLISVLLLSGLTWRR
IL-2Rα_Bear EYQIAVAGCVLLLISVLLLSGLTWQR
IL-2Rα_Hamster EHQIAVASCVFLLISILLLSGLTWQR
IL-2Rα_Pig QYQLAVAGCVLLLLSILLLSGLTWQR
IL-2Rα_Bat EYQIAVAGCILLLISIFLLSVLTWQR
IL-2Rα_Cattle EYQIAVAGCTLLLASILLLSCLTWQR
IL-2Rα_Sheep EYQIAVAGCILLLSSILLLSCLTWQR
IL-2Rα_Whale EYQIAVAGCILLLISIFLLTGLTWQQ
IL-2Rα_Rabbit EYQIAVAGCVFLLISVLLLSALSWRR
IL-2Rα_Sifakas EYQVAVAGCVLLLFTVLLLSGLTWQR
IL-2Rα_Beaver PTSSGVAGCVFLLISVFVLSGLTWQR
250 260 270
IL-2Rβ_Mouse WLRYLLLVLG.CFSGFFSCVYILVKC
IL-2Rβ_Human WLGHLLVGLS.GAFGFIILVYLLINC
IL-2Rβ_Bonobo WLGHLLVGLS.GAFGFIILVYLLINC
IL-2Rβ_Gelada WLGHLLVGLS.GAFGFIILVYLLINC
IL-2Rβ_Rat WLRCLLLVLG.CFFGFLSCVCVLVKC
IL-2Rβ_Horse WWGHTLVGLTGGSLGFIFLVYLLVNC
IL-2Rβ_Cat WLGHIIMGIS.SAFGFIISVYLLANC
IL-2Rβ_Dog WFGHIIMGIT.GAFGFIILVYLLVNC
IL-2Rβ_Monkey WLGHFLVGLS.GAFGFIILVYLLINC
IL-2Rβ_Fox WFGHIIMGVT.GAFGFIILVYLLVNC
IL-2Rβ_Alpaca WLSHVILGLG.SAFGFVFLVYFLANC
IL-2Rβ_Camel WLSHVILGLG.SAFGFVFLVYFLANC
IL-2Rβ_Seal WFGSIITGVS.SACGFIVLVCLLANC
IL-2Rβ_Tiger WLGHITMGIS.SAFGFIISVCLLANC
IL-2Rβ_Bear WFGHVIMGVS.SALGFIILVYLLANC
IL-2Rβ_Pig WSGHIILGLVFAAVSFVFLVYFLAKC
IL-2Rβ_Bat SLGHILVGLC.GALGFLVVVSLLVNC
IL-2Rβ_Cattle WLNHIFLGVG.SFFGLVLLVYFLGSF
IL-2Rβ_Sheep WLVPIFLGVG.SFFGLVLLVYFLGSF
IL-2Rβ_Whale SMGHIVLGLC.CTFGFVLLVYLLTSF
IL-2Rβ_Sifakas GLDYILMGLSAGAVGFIILVYLLVNS
IL-2Rβ_Beaver WFGHFTLGLG.VVVSFLISVYFLAKF
210 220
IL-15R_Mouse TKVAISTSVL..LVGAGVVMAFLAWY
IL-15R_Human TTVAISTSTV.LLCGLSAV.SLLACY
IL-15R_Bonobo TTVAISTSTV.LLCGLSAV.SLLACY
IL-15R_Gelada TTVAISASTV.LLCGLSAV.SLLACY
IL-15R_Rat TKVAIPTSVLLVAAGIGLV..FLTRY
IL-15R_Horse VTAAVSIPVALL.FG.CAV.FLLVRC
IL-15R_Cat VTAAIVMPVAVL.CVVCVG..LLVCY
IL-15R_Dog VTAAIFTPVTVL.CVVCVV.FLLVCY
IL-15R_Monkey LSVAISTSTV.LLCGLSAV.SLLACY
IL-15R_Fox VTAAIFTPVTVL.CVVCVV.FLLVCY
IL-15R_Alpaca VTAAVSVLVAVL.GAVCVG.LLLVCY
IL-15R_Camel VTAAVSVLVAVL.GAVCVG.LLLVCY
IL-15R_Seal VTVAIFTPVAVL.GVVCVV.LLLVCY
IL-15R_Tiger VTAAIVMPVTVL.CVVCVG..LLVCY
IL-15R_Bear VTAAIFTPVAVL.SVMCVV.LLLVCY
IL-15R_Hamster TKVAISTSVLTLLVGAGVV.ALLVCY
IL-15R_Pig VAVAVSTPCALLLCGACVL.LLVVHF
IL-15R_Cattle VTAAVST.VTVL.FVVCLV.FLLGRC
IL-15R_Sheep VTAAVSIPVAVV.FAVCLV.FLLVRC
IL-15R_Whale VTAAISTSVAVL.CGVCVV.CLLVRY
IL-15R_Rabbit VTVAISTSVSLL.GVVCVG.FLLAYC
IL-15R_Sifakas ATVTISTAAVLL.CVLSGA..LLACC
IL-15R_Beaver AIVTISTSVSTF.CTVLVM.LLLLWC
240 250
IL-21R_Mouse DPHMLLLLAVLII.VLV.F.MGLKIH
IL-21R_Human NPHLLLLLLLVIVFIPA.F.WSLKTH
IL-21R_Bonobo NPHLLLLLLLVIVFIPA.F.WSLKTH
IL-21R_Gelada NPHLLLLLLLVIVFIPA.F.WSLKTH
IL-21R_Rat DPHVLLLLAVLIL.VLV.F.MGLKNH
IL-21R_Horse HSDLLYLLLVLVPPILV.F.LGLKIH
IL-21R_Cat HIYLLYIFLVIV..VIV.F.LGLKID
IL-21R_Dog HTYLLYLLLVIILPILV.F.LGLKIH
IL-21R_Monkey NPHLLLLLLLVIVFIPA.F.WSLKTH
IL-21R_Fox HTYLLYLLLVIILPILV.F.LGLKIH
IL-21R_Alpaca YPHLLPIL.VLVSPILV.F.LGLKIH
IL-21R_Camel YPHLLPIL.VLVSPILV.F.LGLKIH
IL-21R_Seal HTYLLYLLLVIILPILV.F.LGLKIH
IL-21R_Tiger HIYLLYIFLVIV..VIV.F.LGLKID
IL-21R_Bear RTYLLYLLLVIILPILV.F.LGLKIH
IL-21R_Hamster DPHLLLLLLIVSAPVLI.I.MGLKNH
IL-21R_Pig YPHLLLIL.ILVCPVLV.F.LGLKIH
IL-21R_Cattle YLHLVPIPLILVCLIFV.F.LGLKVP
IL-21R_Sheep YLHLVPILLILVCLIFV.F.LGLKVP
IL-21R_Whale YPQLL.LILVIVCLIPV.I.LGLKIH
IL-21R_Rabbit DPHLLLLLLVLVAPILV.F.LGLKIH
IL-21R_Sifakas DPHLLLLLLLLIIVCSL.VGWSLKSR
IL-21R_Beaver DPLLLLLPLVFVMPILIIF.LGLKIH
340 350 360
IL-5R_Mouse LVEWHLIVLPTAACFVLLIFSLICRV
IL-5R_Human LREWFVIVIMATICFILLILSLICKI
IL-5R_Monkey LRQWFLIVLMATICFILLILLLICRI
IL-5R_Chick LIVWSLTVLCVSTCSTVTLIAIVCKS

**Fig. S2. Sequence alignment of TMDs across species of the γc family receptors and IL-5R from the βc family.**

The TMD sequences of γc family receptors and the βc family member IL-5R are aligned using COBALT (https://www.ncbi.nlm.nih.gov/tools/cobalt/cobalt.cgi) and colored according to identity percentage. Residues with high similarity across species are shown in red. The 100% conserved residues are shown in white with red background. Transmembrane regions are highlighted in blue box.

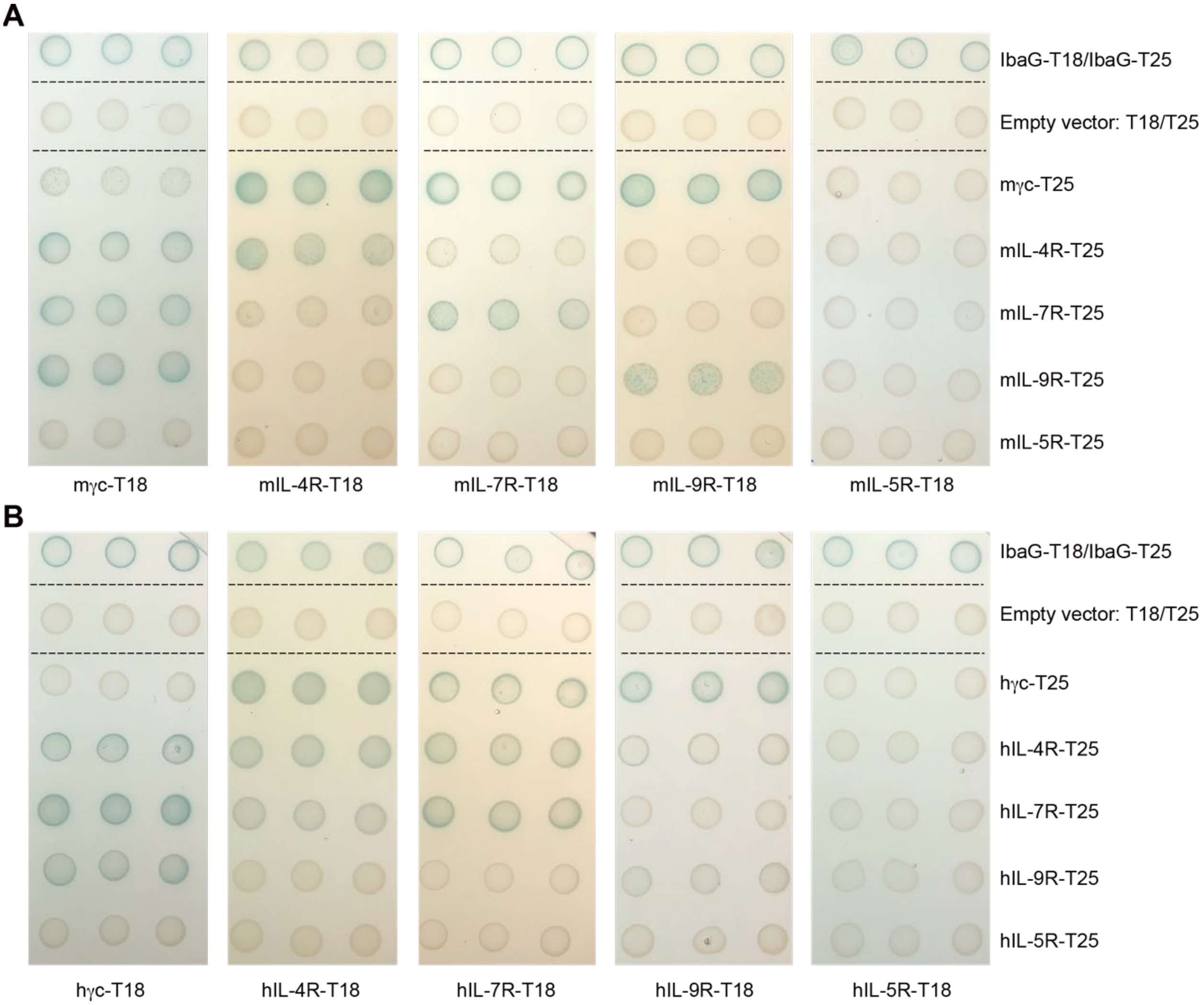


**Fig. S3. Additional results of BACTH analysis of heterotypic and homotypic TMD interactions.**

(**A**) BACTH analysis of TMD interactions for representative γc family receptors including γc, IL-4R, IL-7R and IL-9R as well as the βc family receptor IL-5R from mouse.

(**B**) Same as in (A) for TMD sequences from human.

Three colonies were tested for each TMD-TMD combination. IbaG fused with T18 and T25 (IbaG-T18/IbaG-T25) was used as the positive control as previously published (*39*). Co-expression of empty vectors of T18 and T25 was used as the negative control. Blue colonies indicate TMD-TMD association in the bacteria inner membrane.

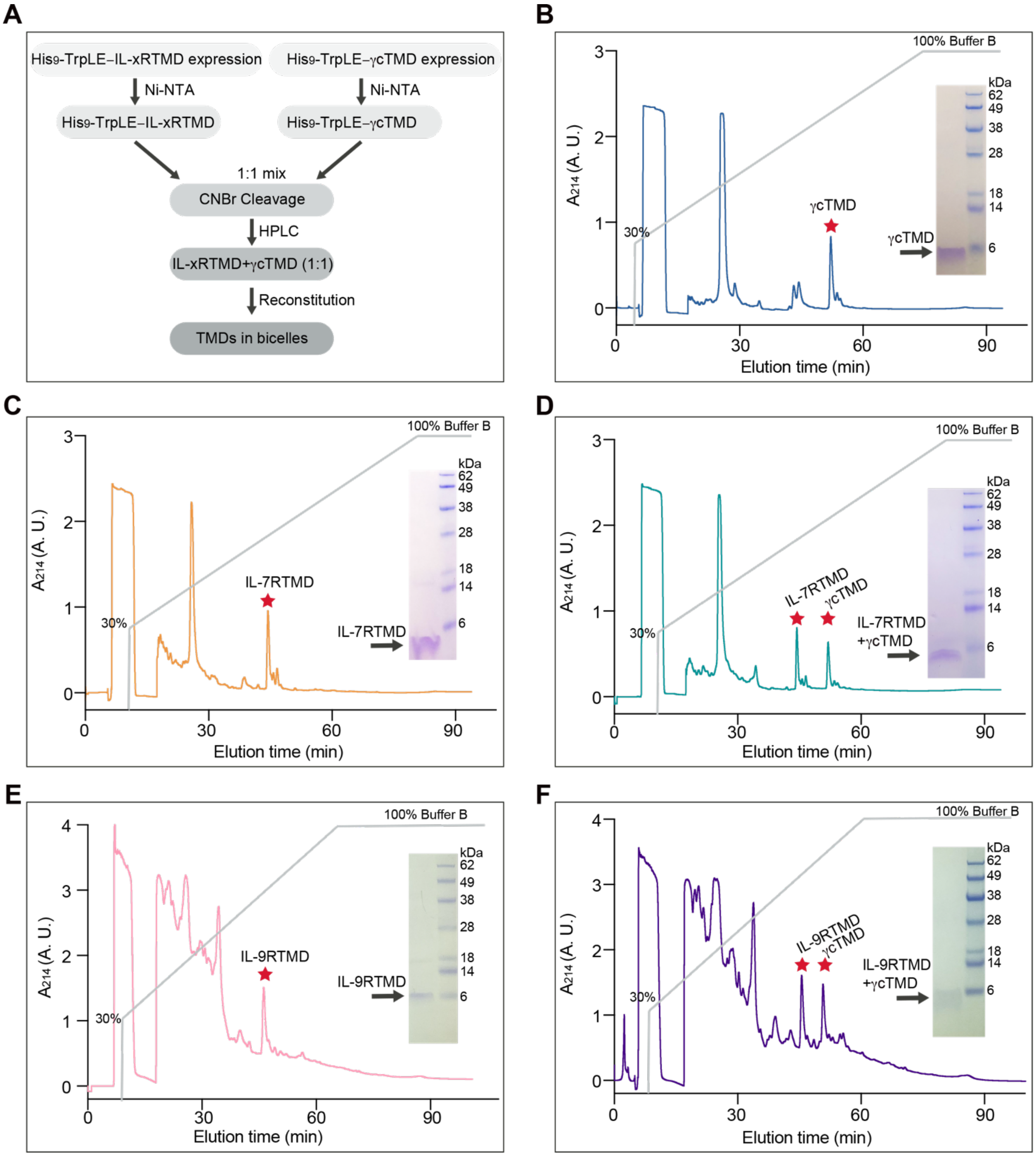


**Fig. S4. Purification of γcTMD, IL-7RTMD, IL9R-TMD and mixed TMDs**

(**A**) Overview of the steps for mixed TMDs expression and purification (see Supplementary Materials and Methods for detailed description).

(**B**) Reverse phase HPLC purification of γcTMD from CNBr-cleaved TrpLE-γcTMD fusion protein on Zorbax SB-C3 column with a gradient from 95% dH2O, 5% isopropanol (IPA), 0.1% trifluoroacetic acid (TFA) (buffer A) to 75% IPA, 25% acetonitrile, 0.1% TFA (buffer B).

(**C**) Same as in (B) for IL-7RTMD.

(**D**) Same as in (B) for mixed IL-7RTMD with γcTMD.

(**E**) Same as in (B) for IL-9RTMD.

(**F**) Same as in (B) for mixed IL-9RTMD with γcTMD.

The purity of the TMDs was verified by SDS-PAGE.

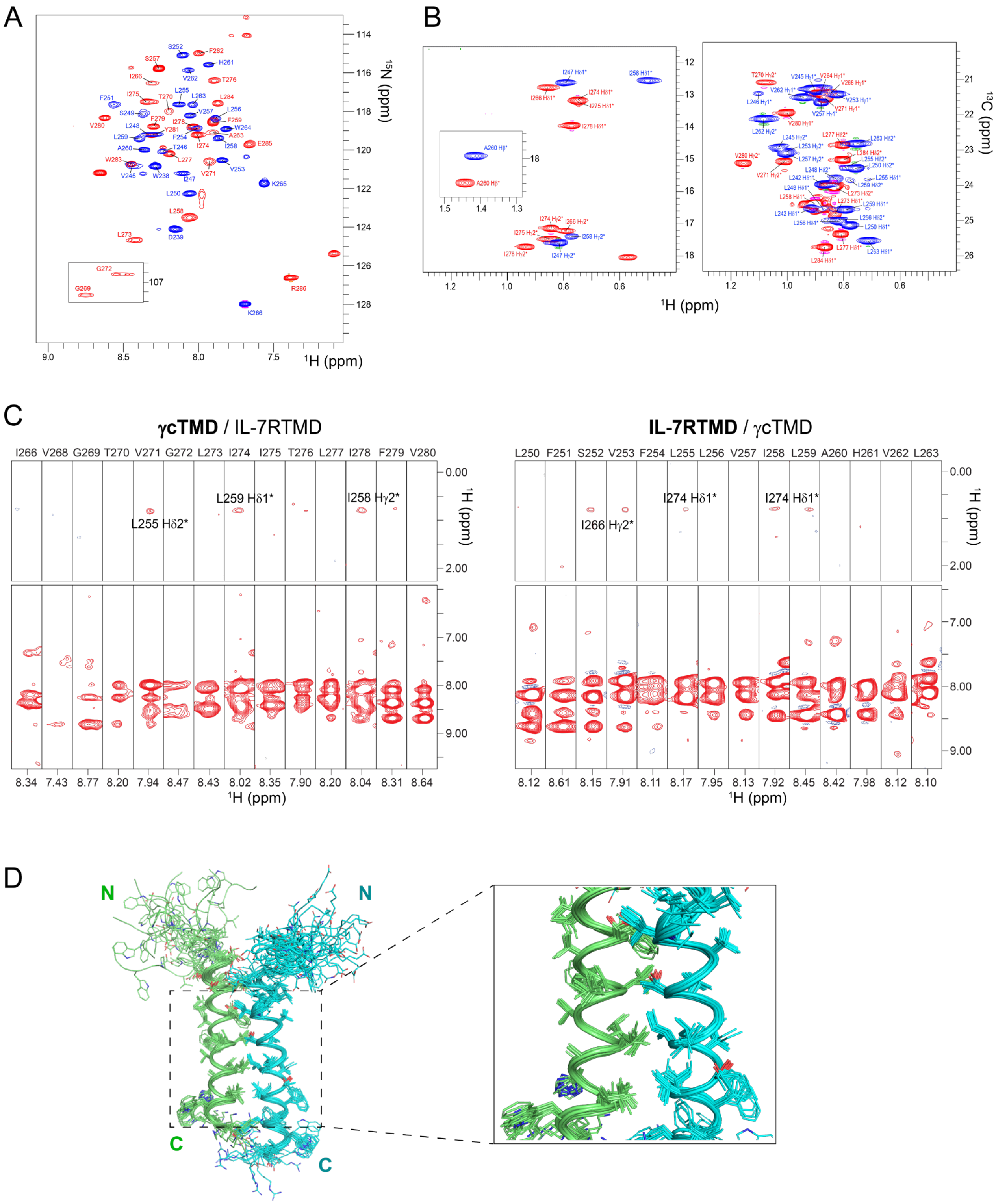


**Fig. S5. NMR characterization of the γcTMD / IL-7RTMD complex.**

(**A**) Overlay of $^{1}H$-$^{15}N$ TROSY-HSQC spectra of ($^{15}N$, $^{2}H$) γcTMD in complex with $^{13}C$ IL-7RTMD (red) and the reciprocal sample of ($^{15}N$, $^{2}H$) IL-7RTMD in complex with $^{13}C$ γcTMD (blue), recorded at 303K at $^{1}H$ frequency of 900 MHz. The glycine peaks are folded.
(**B**) Overlay of $^{1}H$-$^{13}C$ HSQC spectra (methyl groups) of $^{13}C$ γcTMD in complex with ($^{15}N$, $^{2}H$) IL-7RTMD (red) and $^{13}C$ IL-7RTMD in complex with ($^{15}N$, $^{2}H$) γcTMD (blue).
(**C**) Strips from 3D $^{15}N$-edited NOESY-TROSY spectra of ($^{15}N$, $^{2}H$) γcTMD / ($^{13}C$) IL-7RTMD (left) and ($^{15}N$, $^{2}H$) IL-7RTMD / ($^{13}C$) γcTMD (right) showing inter-chain NOEs between the amide protons of $^{15}N$ labeled and fully deuterated chain and the methyl protons of the $^{13}C$ labeled chain. The spectra were recorded at 303K at $^{1}H$ frequency of 900 MHz.
(**D**) Ensemble of 15 lowest energy structures from 100 structures calculated using all NMR-derived restraints in Table S1. Protons are not displayed for clarity. Residues included in the superposition are 262-284 of γcTMD and 245-265 of IL-7RTMD.

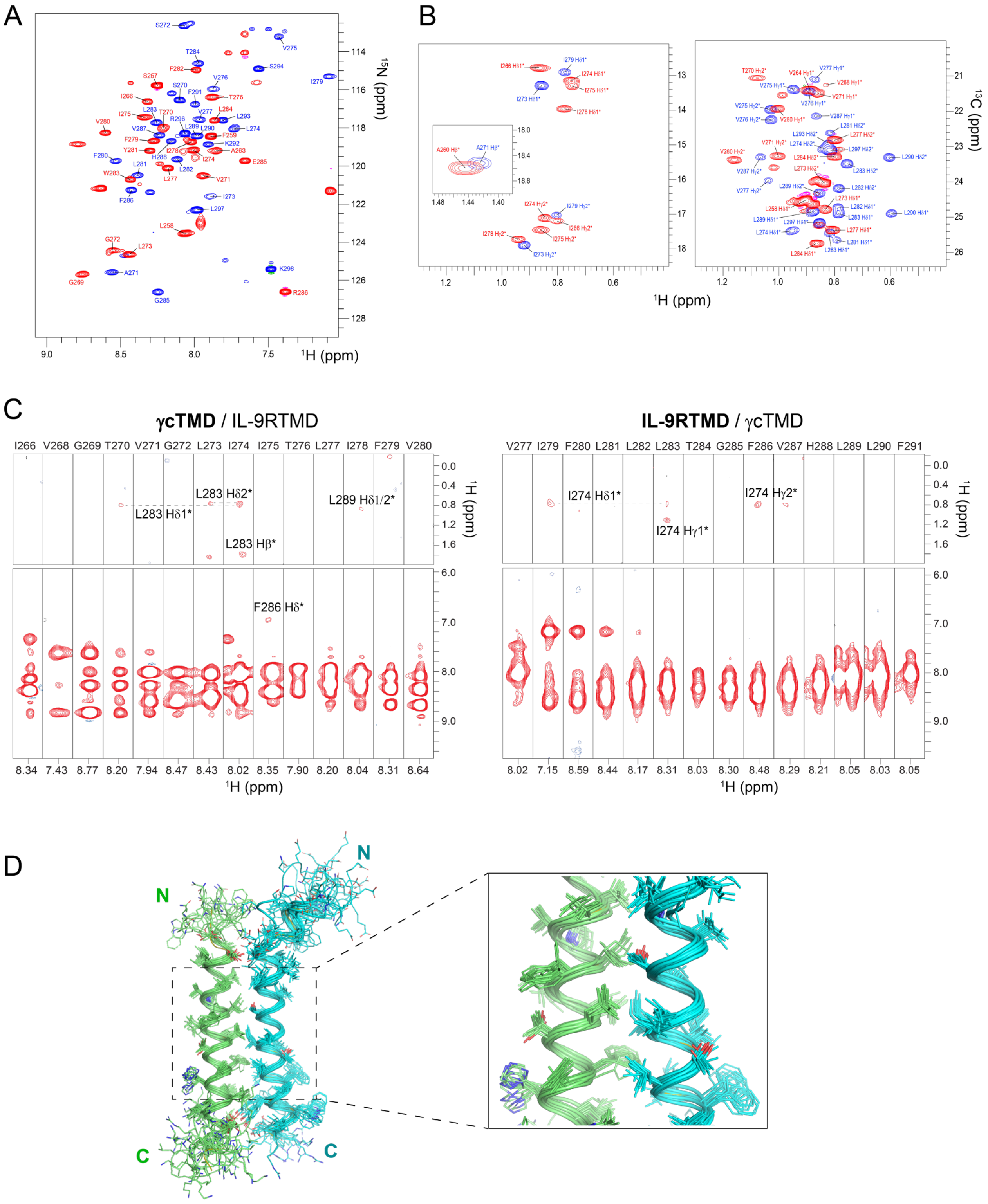


**Fig. S6. NMR characterization of the γcTMD / IL-9RTMD complex.**

(**A**) Overlay of $^1$H-$^{15}$N TROSY-HSQC spectra of ($^{15}$N, $^2$H) γcTMD in complex with $^{13}$C IL-9RTMD (red) and the reciprocal sample of ($^{15}$N, $^2$H) IL-9RTMD in complex with $^{13}$C γcTMD (blue), recorded at 303K at $^1$H frequency of 800 MHz. The glycine peaks are folded.

(**B**) Overlay of $^1$H-$^{13}$C HSQC spectra (methyl groups) of $^{13}$C γcTMD in complex with ($^{15}$N, $^2$H) IL-7RTMD (red) and $^{13}$C IL-7RTMD in complex with ($^{15}$N, $^2$H) γcTMD (blue).

(**C**) Strips from 3D $^{15}$N-edited NOESY-TROSY spectra of ($^{15}$N, $^2$H) γcTMD / ($^{13}$C) IL-9RTMD (left) and ($^{15}$N, $^2$H) IL-9RTMD / ($^{13}$C) γcTMD (right) showing inter-chain NOEs between the amide protons of $^{15}$N labeled and fully deuterated chain and the methyl protons of the $^{13}$C labeled chain. The spectra were recorded at 303K at $^1$H frequency of 800 MHz.

(**D**) Ensemble of 15 lowest energy structures from 100 structures calculated using all NMR-derived restraints in Table S1. Protons are not displayed for clarity. Residues included in the superposition are 262-284 of γcTMD and 271-293 of IL-9RTMD.

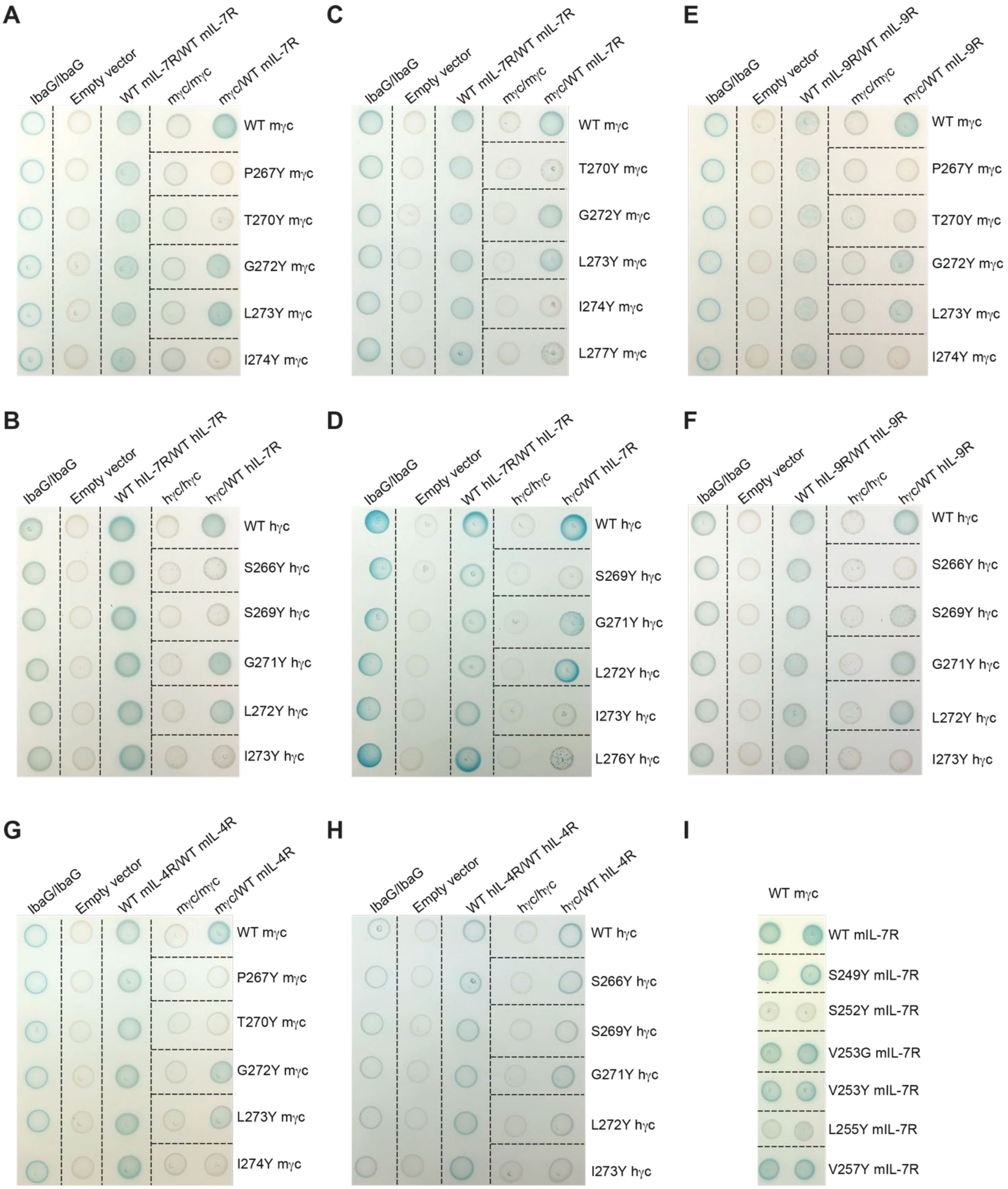


**Fig. S7. Additional results of site-directed mutagenesis and effect on TMD interaction.** (**A-D**) Raw images of BACTH analysis of the effect of γcTMD mutations (P267Y, T270Y, G272Y, L273Y, I274Y, and L277Y in mouse γcTMD, and the corresponding mutations S266Y,

S269Y, G271Y, L272Y, I273Y, and L276Y in human) on γcTMD association with mouse (**A** and **C**) and human (**B** and **D**) WT IL-7RTMD.

(**E** and **F**) Same as in (A) and (B) for WT IL-9RTMD.

(**G** and **H**) Same as in (A) and (B) for WT IL-4RTMD.

(**I**) Raw image of BACTH analysis of the effect of mouse IL-7RTMD mutations on association with WT mouse γcTMD.

IbaG fused with T18 and T25 (IbaG-T18/IbaG-T25) was used as the positive control as previously published(*39*). Co-expression of empty vectors of T18 and T25 was used as the negative control. Blue colonies indicate TMD-TMD association in the bacteria inner membrane.

**Table S1. NMR structure calculation and refinement statistics**

| NMR distance and dihedral constraints [a] | γcTMD / IL-7RTMD | γcTMD / IL-9RTMD |
|---|---|---|
| Distance constraints from NOE | 248 | 230 |
| Short-range intramolecular ($\|i-j\| \leq 4$) | 226 | 214 |
| Long-range intramolecular ($\|i-j\| \geq 5$) | 0 | 0 |
| Intermolecular | 22 | 16 |
| Total dihedral angle restraints [b] | 94 | 100 |
| $\phi$ (TALOS) | 47 | 50 |
| $\psi$ (TALOS) | 47 | 50 |
| **Structure statistics** [c] | | |
| Violations (mean ± s.d.) | | |
| Distance constraints (Å) | 0.079 ± 0.004 | 0.054 ± 0.004 |
| Dihedral angle constraints (°) | 0.210 ± 0.042 | 0.335 ± 0.024 |
| Deviations from idealized geometry | | |
| Bond lengths (Å) | 0.006 ± 0.000 | 0.006 ± 0.000 |
| Bond angles (°) | 0.284 ± 0.037 | 0.687 ± 0.012 |
| Impropers (°) | 0.434 ± 0.022 | 0.450 ± 0.020 |
| Average pairwise r.m.s. deviation (Å) [d] | | |
| Heavy | 1.234 | 1.281 |
| Backbone | 0.645 | 0.711 |

[a] The numbers of restraints for the γcTMD/IL-7RTMD complex are summed over residues 257-285 of γcTMD and residues 246-266 of IL-7RTMD. The numbers of restraints for the γcTMD/IL-9RTMD complex are summed over residues 258-285 of γcTMD and residues 271-297 of IL-9RTMD.

[b] Backbone $\phi$ and $\psi$ restraints and their respective uncertainties were obtained from the "GOOD" dihedrals generated by the TALOS+ program (*48*) based on the backbone chemical shift values.

[c] Statistics are calculated and averaged over an ensemble of the 15 lowest energy structures out of 100 calculated structures.

[d] The precision of the atomic coordinates is defined as the average r.m.s. difference between the 15 final structures and their mean coordinates. The calculation only includes the ordered regions of the protein: γcTMD residues 262-284 and IL-7RTMD residues 245-265 of the γcTMD/IL-7RTMD complex; γcTMD residues 262-284 and IL-9RTMD residues 271-293 of the γcTMD/IL-9RTMD complex.

**Table S2. Cell lines, plasmids and reagents used in this study**

| Reagent or Resource | Source | Identifier |
|---|---|---|
| **Antibodies** | | |
| Phospho-STAT5 (Tyr694) (C11C5) | Cell Signaling Technology | #9359 |
| Beta-actin | Cell signaling | #4970S |
| Anti-rabbit IgG-HRP linked Ab | Cell signaling | #7074S |
| **Bacterial Strains and mammalian cells** | | |
| E. coli BL21(DE3) | New England Biolabs | Cat# C2527 |
| E. coli DH5-alpha | New England Biolabs | Cat# C2987 |
| E. coli C43 (DE3) | Sigma-Aldrich | CMC0019 |
| BTH101 | Euromedex | EUB001 |
| HEK293T | ATCC | CRL-3216; RRID: CVCL_0063 |
| BaF3 | AcceGen Biotech | ABC-TC060S |
| **Chemicals, Peptides, and Recombinant Proteins** | | |
| Cyanogen Bromide | Sigma-Aldrich | Cat# C91492 |
| Deuterium oxide | Cambridge Isotope Laboratories (CIL) | DLM-4-99.8-1000 |
| $^{13}$C D-Glucose | CIL | CLM-13965 |
| D1,2,3,4,5,6,7 D-Glucose | CIL | DLM-20621 |
| $^{15}$N ammonium chloride | CIL | NLM-467-5 |
| Kanamycin monosulfate | Sigma-Aldrich | Cat# BP861 |
| Ampicillin, Sodium Salt | Sigma-Aldrich | Cat# 171254 |
| isopropyl b-D-thiogalatopyranoside (IPTG) | Sigma-Aldrich | Cat# I5502 |
| DMPC Lipid | Avanti Polar Lipids | Cat# 850345 |
| DH6PC Detergent | Avanti Polar Lipids | Cat# 850305 |
| Deuterated d54-DMPC lipid | Cortecnet | CD5012P025 |
| Deuterated d22-DH6PC Detergent | Cortecnet | CD5010P025 |
| Recombinant mouse IL-3 | SIGMA-ALDRICH | #SRP3208-10UG |

| | | |
|---|---|---|
| Recombinant human IL-7 | Thermo Fisher Scientific | #PHC0075 |
| TEV enzyme | James Chou lab | N/A |
| DAPI | Abcam | #ab228549 |
| Polybrene | Sigma-Aldrich | #H9268 |
| Retro-X™ Concentrator | Takara | #631455 |
| RPMI 1640 medium | Thermo Fisher Scientific | # 11875119 |
| DMEM | Gibco | #10566-016 |
| Fetal bovine serum | Thermo Fisher Scientific | # 10082147 |
| Penicillin-Streptomycin | Thermo Fisher Scientific | #15140122 |
| DPBS | Gibco | #14190-144 |
| Trypsin-EDTA | Gibco | # 25200056 |
| Lipofectamine™ 3000 | Thermo Fisher Scientific | # L3000008 |
| Opti-MEM | Gibco | #31985-062 |
| X-Gal | AdipoGen | #AG-CC1-0003 |
| **Vectors and plasmids** | | |
| pKNT25 | Euromedex | EUP-25N |
| pUT18 | Euromedex | EUP-18N |
| IbaG-pKNT25 | Abdelrahim Zoued | N/A |
| IbaG- pUT18 | Abdelrahim Zoued | N/A |
| OmpA-mIL-7RTD-pKNT25 | This paper | N/A |
| OmpA-mIL-7RTMD-pUT18 | This paper | N/A |
| OmpA-hIL-7RTMD-pKNT25 | This paper | N/A |
| OmpA-hIL-7RTMD-pUT18 | This paper | N/A |
| OmpA-mγcTMD-pKNT25 | This paper | N/A |
| OmpA-mγcTMD-pUT18 | This paper | N/A |
| OmpA-hγcTMD-pKNT25 | This paper | N/A |

| | | |
|---|---|---|
| OmpA-hγcTMD-UT18 | This paper | N/A |
| OmpA-mIL-4RTMD-pKNT25 | This paper | N/A |
| OmpA-mIL-4RTMD-pUT18 | This paper | N/A |
| OmpA-hIL-4RTMD-pKNT25 | This paper | N/A |
| OmpA-hIL-4RTMD-pUT18 | This paper | N/A |
| OmpA-mIL-9RTMD-pKNT25 | This paper | N/A |
| OmpA-mIL-9RTMD-pUT18 | This paper | N/A |
| OmpA-hIL-9RTMD-pKNT25 | This paper | N/A |
| OmpA-hIL-9RTMD-pUT18 | This paper | N/A |
| OmpA-mIL-5RTMD-pKNT25 | This paper | N/A |
| OmpA-mIL-5RTMD-pUT18 | This paper | N/A |
| OmpA-hIL-5RTMD-pKNT25 | This paper | N/A |
| OmpA-hIL-5RTMD-pUT18 | This paper | N/A |
| OmpA-P267Y mγcTMD-pKNT25 | This paper | N/A |
| OmpA-P267Y mγcTMD-pUT18 | This paper | N/A |
| OmpA-T270Y mγcTMD-pKNT25 | This paper | N/A |
| OmpA-T270Y mγcTMD-pUT18 | This paper | N/A |
| OmpA-G272Y mγcTMD-pKNT25 | This paper | N/A |
| OmpA-G272Y mγcTMD-pUT18 | This paper | N/A |
| OmpA-L273Y mγcTMD-pKNT25 | This paper | N/A |
| OmpA-L273Y mγcTMD-pUT18 | This paper | N/A |
| OmpA-I274Y mγcTMD-pKNT25 | This paper | N/A |
| OmpA-I274Y mγcTMD-pUT18 | This paper | N/A |
| OmpA-L277Y mγcTMD-pKNT25 | This paper | N/A |
| OmpA-L277Y mγcTMD-pUT18 | This paper | N/A |
| OmpA-S266Y hγcTMD-pKNT25 | This paper | N/A |
| OmpA-S266Y hγcTMD-pUT18 | This paper | N/A |
| OmpA-S269Y hγcTMD-pKNT25 | This paper | N/A |
| OmpA-S269Y hγcTMD-pUT18 | This paper | N/A |

| | | |
|---|---|---|
| OmpA-G271Y hγcTMD-pKNT25 | This paper | N/A |
| OmpA-G271Y hγcTMD-pUT18 | This paper | N/A |
| OmpA-L272Y hγcTMD-pKNT25 | This paper | N/A |
| OmpA-L272Y hγcTMD-pUT18 | This paper | N/A |
| OmpA-I273Y hγcTMD-pKNT25 | This paper | N/A |
| OmpA-I273Y hγcTMD-UT18 | This paper | N/A |
| OmpA-L276Y hγcTMD-pKNT25 | This paper | N/A |
| OmpA-L276Y hγcTMD-UT18 | This paper | N/A |
| OmpA-S249Y mIL-7RTMD-pKNT25 | This paper | N/A |
| OmpA- S249Y mIL-7RTMD-pUT18 | This paper | N/A |
| OmpA-S252Y mIL-7RTMD-pKNT25 | This paper | N/A |
| OmpA- S252Y mIL-7RTMD-pUT18 | This paper | N/A |
| OmpA-V253G mIL-7RTMD-pKNT25 | This paper | N/A |
| OmpA- V253G mIL-7RTMD-pUT18 | This paper | N/A |
| OmpA-V253Y mIL-7RTMD-pKNT25 | This paper | N/A |
| OmpA- V253Y mIL-7RTMD-pUT18 | This paper | N/A |
| OmpA-L255Y mIL-7RTMD-pKNT25 | This paper | N/A |
| OmpA- L255Y mIL-7RTMD-pUT18 | This paper | N/A |
| OmpA-V257Y mIL-7RTMD-pKNT25 | This paper | N/A |
| OmpA- V257Y mIL-7RTMD-pUT18 | This paper | N/A |
| MSCV-hIL-7R-EGFP | This paper | N/A |
| MSCV-hIL-7R TEV-EGFP | This paper | N/A |
| MSCV-hIL-7R V253Y-EGFP | This paper | N/A |
| MSCV-hIL-7R L255Y-EGFP | This paper | N/A |
| MSCV-hIL-7R-ΔECD-EGFP | This paper | N/A |
| MSCV- hγc-mCherry | This paper | N/A |
| MSCV- hγc TEV-mCherry | This paper | N/A |
| MSCV- hγc G271Y-mCherry | This paper | N/A |
| MSCV- hγc I273Y-mCherry | This paper | N/A |

| | | |
|---|---|---|
| MSCV- hγc-ΔECD -mCherry | This paper | N/A |
| pMM-LR6 -His-TrpLE-mγcTMD | This paper | N/A |
| pMM-LR6 -His-TrpLE-mIL-7RTMD | This paper | N/A |
| pMM-LR6 -His-TrpLE-mIL-9RTMD | This paper | N/A |
